# Dimerization mechanism and structural features of human LI-cadherin

**DOI:** 10.1101/2020.09.18.291195

**Authors:** Anna Yui, Jose M. M. Caaveiro, Daisuke Kuroda, Makoto Nakakido, Satoru Nagatoishi, Shuichiro Goda, Takahiro Maruno, Susumu Uchiyama, Kouhei Tsumoto

## Abstract

LI-cadherin is a member of the cadherin superfamily, which encompasses a group of Ca^2+^-dependent cell adhesion proteins. The expression of LI-cadherin is observed on various types of cells in the human body, such as normal small intestine and colon cells, and gastric cancer cells. Because its expression is not observed on normal gastric cells, LI-cadherin is a promising target for gastric cancer imaging. However, since the cell adhesion mechanism of LI-cadherin has remained unknown, rational design of therapeutic molecules targeting this cadherin has been hampered. Here, we have studied the homodimerization mechanism of LI-cadherin. We report the crystal structure of the LI-cadherin EC1-4 homodimer. The EC1-4 homodimer exhibited a unique architecture different from that of other cadherins reported so far. The crystal structure also revealed that LI-cadherin possesses a noncanonical calcium ion-free linker between EC2 and EC3. Various biochemical techniques and molecular dynamics (MD) simulations were employed to elucidate the mechanism of homodimerization. We also showed that the formation of the homodimer observed by the crystal structure is necessary for LI-cadherin-dependent cell adhesion by performing cell aggregation assay.

## Introduction

Cadherins are a family of glycoproteins responsible for calcium ion-dependent cell adhesion (1). There are more than 100 types of cadherins in humans and many of them are responsible not only for cell adhesion but also involved in tumorigenesis (2). Human liver intestine-cadherin (LI-cadherin) is a nonclassical cadherin composed of an ectodomain consisting of seven extracellular cadherin (EC) repeats, a single transmembrane domain, and a short cytoplasmic domain (3). Previous studies have reported the expression of LI-cadherin on various types of cells, such as normal intestine cells, intestinal metaplasia, colorectal cancer cells, and lymph node metastatic gastric cancer cells (4, 5).

Because human LI-cadherin is expressed on gastric cancer cells but not on normal stomach tissues, LI-cadherin has been proposed as a target for imaging metastatic gastric cancer (6). Previous studies have reported that LI-cadherin works not only as a calcium ion-dependent cell adhesion molecule as other cadherins do (7), but also shown that trans-dimerization of LI-cadherin is necessary for water transport in normal intestinal cells (8). Sequence alignment of mouse LI-, E-, N-, and P-cadherins has revealed the sequence homology between EC1-2 of LI-cadherin and EC1-2 of E-, N-, and P-cadherins, as well as between EC3-7 of LI-cadherin and EC1-5 of classical cadherins (9). From the sequence similarity and the proposed absence of calcium ion-binding motifs (10, 11) between EC2 and EC3 repeats, there is speculation that LI-cadherin has evolved from the same five-repeat precursor as that of classical cadherins (9).

However, LI-cadherin is different from classical cadherins in several aspects, such as the number of EC repeats and the length and sequence of the cytoplasmic domain. Classical cadherins possess five EC repeats whereas LI-cadherin displays seven (2). Classical cadherins possess a conserved cytoplasmic domain comprising more than 100 amino acids, whereas that of LI-cadherin is only 20 residues long with little or no sequence homology to that of classical cadherins (7, 12).

The characteristics of LI-cadherin at the molecular level, including the homodimerization mechanism, remain unknown. Homodimerization is the fundamental event in cadherin-mediated cell adhesion as has been shown previously (13, 14). For example, classical cadherins form a homodimer mediated by the interaction between their two N-terminal EC repeats (EC1-2) (10, 15).

In this study, we aimed to characterize LI-cadherin at the molecular level, since the molecular description of the target protein may play a significant role for the rational design of therapeutic approaches. We have extensively validated LI-cadherin to identify the homodimer architecture of LI-cadherin. Here, we report the crystal structure of the homodimer form of human LI-cadherin EC1-4. The crystal structure revealed a dimerization architecture different from that of any other cadherin reported so far. It also showed canonical calcium binding motifs between EC1 and EC2, and between EC3 and EC4, but not between EC2 and EC3. By performing various biochemical and computational analysis based on this crystal structure, we interpreted the characteristics of LI-cadherin molecule. Additionally, we showed that the formation of the EC1-4 homodimer is necessary for LI-cadherin-dependent cell adhesion through cell aggregation assays. Our study revealed possible architectures of LI-cadherin homodimers at the cell surface and suggested the differential role of the two additional EC repeats at the N-terminus compared with classical cadherins.

## Results

### Homodimerization propensity of human LI-cadherin

To investigate the homodimerization mechanism of LI-cadherin, we expressed the entire ectodomain comprising EC1-7 (Table S1) and analyzed the homodimerization propensity using sedimentation velocity analytical ultracentrifugation (SV-AUC). Although formation of homodimer was observed, it was not possible to determine its dissociation constant (*K*_D_) because no concentration-dependence in the weight-average of sedimentation coefficient was discerned, suggesting a very slow dissociation rate (Fig. 2).

Therefore, to understand the homodimerization mechanism of LI-cadherin in more detail, we prepared truncated versions of LI-cadherin containing various numbers of EC repeats and evaluated their homodimerization potency. The constructs were designed based on the sequence homology between LI-cadherin and classical cadherins. We compared the sequence of human LI-cadherin and human classical cadherins (E-, N- and P-cadherins) using EMBOSS Needle (16). As it has been pointed out in a previous study (9), EC1-2 and EC3-7 of human LI-cadherin had an approximately 30% sequence homology with EC1-2 and EC1-5 of human classical cadherins, respectively (Fig. 1 and Tables S2-3). Notably, Trp239 locates at the N-terminal end of LI-cadherin EC3 and because of that it has been suggested that this Trp residue might function as an adhesive element equivalent to that of the conserved residue Trp2 of EC1 of classical cadherins, playing a crucial role in the formation of strand swap-dimer (ss-dimer) (9, 10, 17–19). Considering the degree of sequence homology and that EC1-2 of classical cadherins is the element responsible for homodimerization, we hypothesized that EC1-2 and EC3-4 of LI-cadherin would be responsible for its dimerization. Therefore, we determined the degree of homodimerization of EC1-2 and EC3-4, as well as those of EC1-4 and EC1-5, using SV-AUC (Table S1).

**Figure 1.**
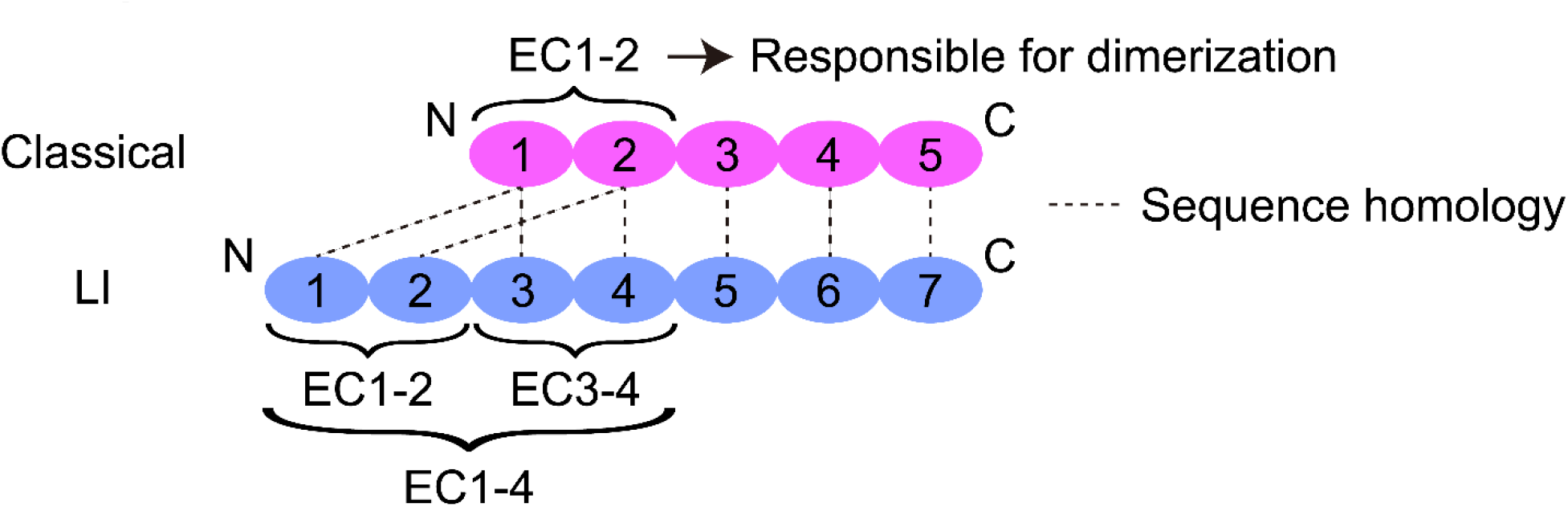
Schematic view of extracellular cadherin (EC) repeats of classical cadherin and LI-cadherin. EC repeats connected by dotted lines indicate sequence homology. EC1-4, EC1-2, and EC3-4 repeats in LI-cadherin, which were employed for the experimental work, are indicated by brackets.

Homodimerization of EC1-2, EC1-4, and EC1-5 was observed, and unlike EC1-7, the weight-average of sedimentation coefficient increased in a concentration-dependent manner. The *K*_D_ values determined were 75.0 μM, 39.8 μM, and 22.8 μM, respectively. In contrast, we did not observe dimerization when employing EC3-4 despite the sequence similarity with EC1-2 of classical cadherins and the presence of Trp239 in EC3, a residue located at the analogous position to that of the key Trp2 residue in EC1 of classical cadherins (Fig. 2). The solution structure of EC3-4 even at the higher concentration was monomeric as determined by small angle X-ray scattering (SAXS), supporting the results of SV-AUC (Fig. S1A~E and Table S4).

**Figure 2.**
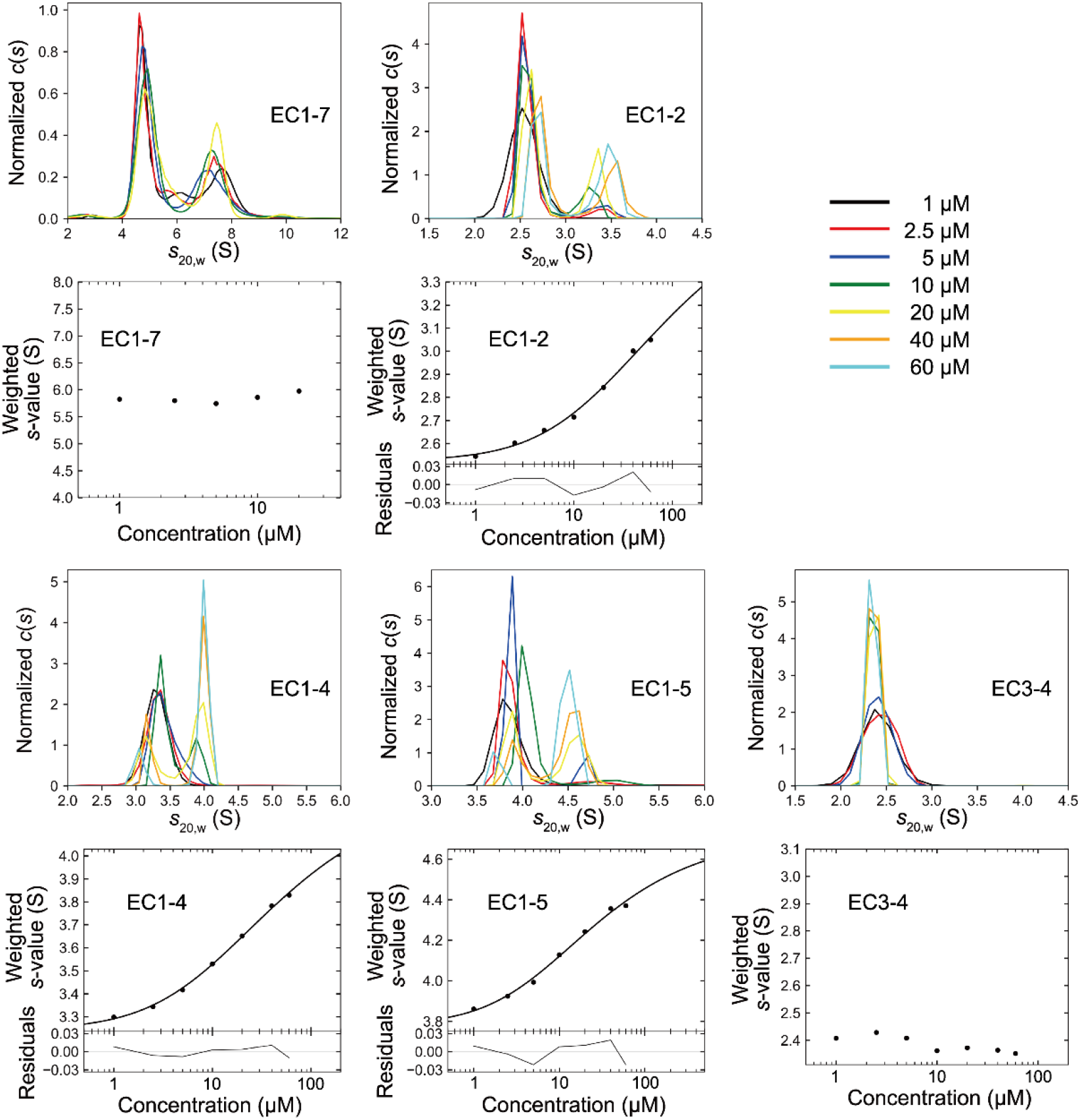
Sedimentation plot of SV-AUC. Dimerization of EC1-7, EC1-2, EC1-4, and EC1-5 were observed. The values of *K*_D_ of homodimerization determined for EC1-2, EC1-4, and EC1-5 were 75.0 μM, 39.8 μM, and 22.8 μM, respectively.

### Crystal structure analysis of EC1-4 homodimer

In order to determine the EC repeats responsible for the homodimerization of LI-cadherin, we determined the crystal structure of EC1-4 expressed in mammalian cells at 2.7 Å resolution (Fig. 3 and Table 1). Each EC repeat was composed of the typical seven β-strands seen in classical cadherins, and three calcium ions bound to each of the linkers connecting EC1 and EC2, and EC3 and EC4 (Fig. 3). We also observed four N-glycans and two N-glycans bound to chain A and B, respectively, as predicted from the amino acid sequence. We could not resolve these N-glycans in their entire length because of their intrinsic flexibility. From the portion resolved, all N-glycans face the opposite side of the dimer interface.

**Figure 3.**
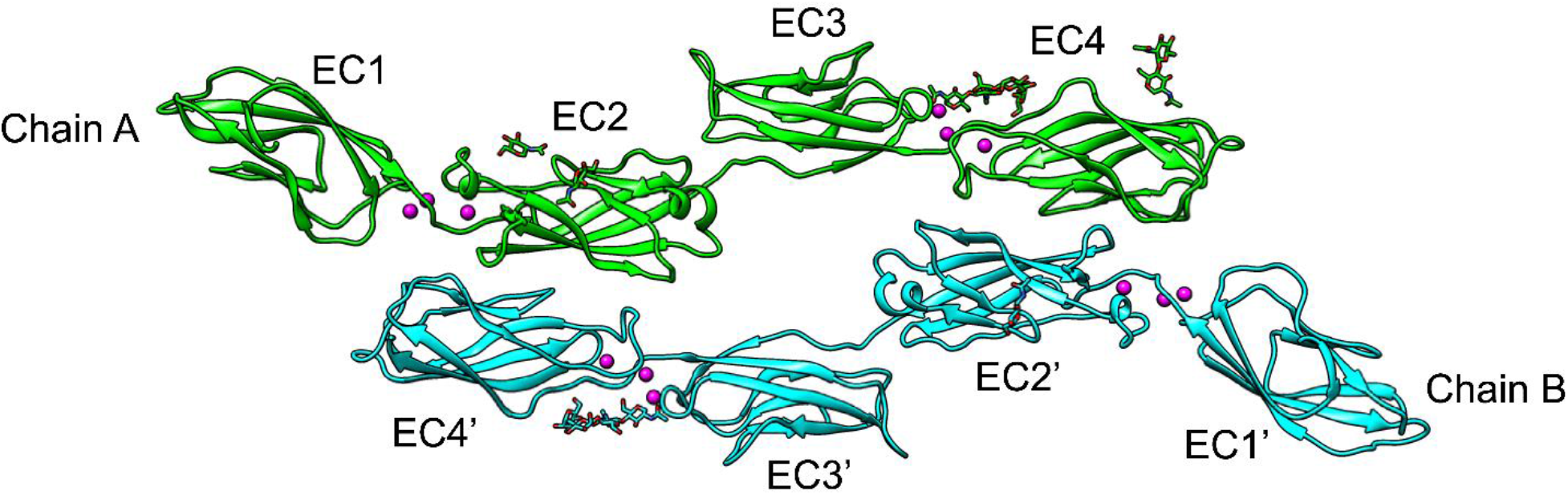
Crystal structure of EC1-4 homodimer. Calcium ions are depicted in magenta. No calcium ions were observed between EC2 and EC3 in either molecule. Four partial N-glycans were modeled in chain A (light green) and two in chain B (cyan) (the amino acid sequence of EC1-4 is given in Table S1).

**Table 1:** Data collection and refinement statistics. Statistical values given in parenthesis refer to the highest resolution bin.

| **Data Collection** | **LI-cadherin (EC1-4)** |
| --- | --- |
| Space Group | P 1 2<sub>1</sub> 1 |
| Unit cell |  |
| a, b, c (Å) | 80.36, 70.84, 134.22 |
| $\alpha$, $\beta$, $\gamma$ (°) | 90.0, 98.7, 90.0 |
| Resolution (Å) | 55.19 - 2.70 (2.85 - 2.70) |
| Wavelength | 1.0000 |
| Observations | 252,491 (37,071) |
| Unique reflections | 41,272 (5,955) |
| *R*<sub>merge</sub> | 0.095 (0.858) |
| *R*<sub>p.i.m.</sub> | 0.041 (0.371) |
| CC<sub>1/2</sub> | 0.998 (0.907) |
| *I* / $\sigma$ (*I*) | 11.6 (1.8) |
| Multiplicity | 6.1 (6.2) |
| Completeness (%) | 99.9 (100.0) |
| **Refinement Statistics** |  |
| Resolution (Å) | 55.19 - 2.70 |
| *R*<sub>work</sub> / *R*<sub>free</sub> (%) | 22.2 / 27.4 |
| No. protein chains | 2 |
| No. atoms |  |
| Protein | 6,834 |
| Ca<sup>2+</sup> | 12 |
| Other atoms | - |
| Water | 33 |
| B-factor (Å<sup>2</sup>) |  |
| Protein | 78.2 |
| Ca<sup>2+</sup> | 69.3 |
| Other atoms | - |
| Water | 62.2 |
| Ramachandran Plot |  |
| Preferred (%) | 85.7 |
| Allowed (%) | 14.3 |
| Outliers (%) | 0 |
| RMSD Bond (Å) | 0.013 |
| RMSD Angle (°) | 1.83 |
| PDB | 7CYM |

Two unique characteristics were observed in the crystal structure of LI-cadherin EC1-4: (i) the existence of a calcium-free linker between EC2 and EC3, and (ii) an unusual homodimerization architecture not described before for cadherins. A previous study had suggested that LI-cadherin lacks a calcium-binding motif between EC2 and EC3 (9) and our crystal structure has confirmed that hypothesis experimentally. Crystal structures of cadherins displaying a calcium-free linker have been reported previously and the biological significance of the calcium-free linker has been discussed (20, 21).

The EC1-4 region of LI-cadherin was assembled as an antiparallel homodimer in which EC2 of one chain interacts with EC4 of the opposite chain. This architecture is different from that of other cadherins, such as classical cadherins, which exhibit two step binding mode (15) and to that of protocadherin γB3, which forms an antiparallel EC1-4 homodimer (14) stabilized by intermolecular interactions in which all the EC repeats participate.

The fact that the affinity of the EC1-5 homodimer is almost twice as high as that of the EC1-4 homodimer suggested the presence of contacts between EC1 and EC5, as can be predicted from the arrangement of EC1 of one chain and EC4 of the other chain in the crystal structure, although this interaction does not seem to be strong. In addition, there was no interaction between EC1-2 of one chain and EC1-2 of the other chain in the crystal structure, suggesting that the architecture of the EC1-2 homodimer detected by SV-AUC should be different from that of the EC1-4 homodimer.

### Calcium-free linker

We investigated the calcium-free linker between EC2 and EC3. Classical cadherins generally adopt a crescent-like shape (18, 22). However, in LI-cadherin, the arch-shape was disrupted at the calcium-free linker region and because of that EC1-4 exhibited unique positioning of EC1-2 with respect to EC3-4.

Generally, three calcium ions bound to the linker between each EC repeat confer rigidity to the structure (11). In fact, a previous study on calcium-free linker of cadherin has shown that the linker showed some flexibility (21). To compare the rigidity of the canonical linker with three calcium ions and the calcium-free linker in LI-cadherin, we performed MD simulations. In addition to the monomeric states, we also used the structure of the EC1-4 homodimer as the initial structure of the simulations. After confirming the convergence of the simulations by calculating RMSD values of Cα atoms (Fig. S2, see Experimental Procedures for the details), we compared the rigidity of the linkers by calculating the RMSD values of Cα atoms of EC1 and EC3, respectively, after superposing those of EC2 alone (Fig. 4A, B). The EC3 in the monomer conformation exhibited the largest RMSD. The RMSD values of EC3 in the homodimer were significantly smaller than those of EC3 in the monomer form. Dihedral angles consisting of Cα atoms of residues at the edge of each EC repeat also indicated that the EC1-4 monomer bends largely at the Ca^2+^-free linker (Fig. S3A-C). These results showed that the calcium-free linker between EC2 and EC3 is more flexible than the canonical linker (Movies 1 and 2).

**Figure 4.**
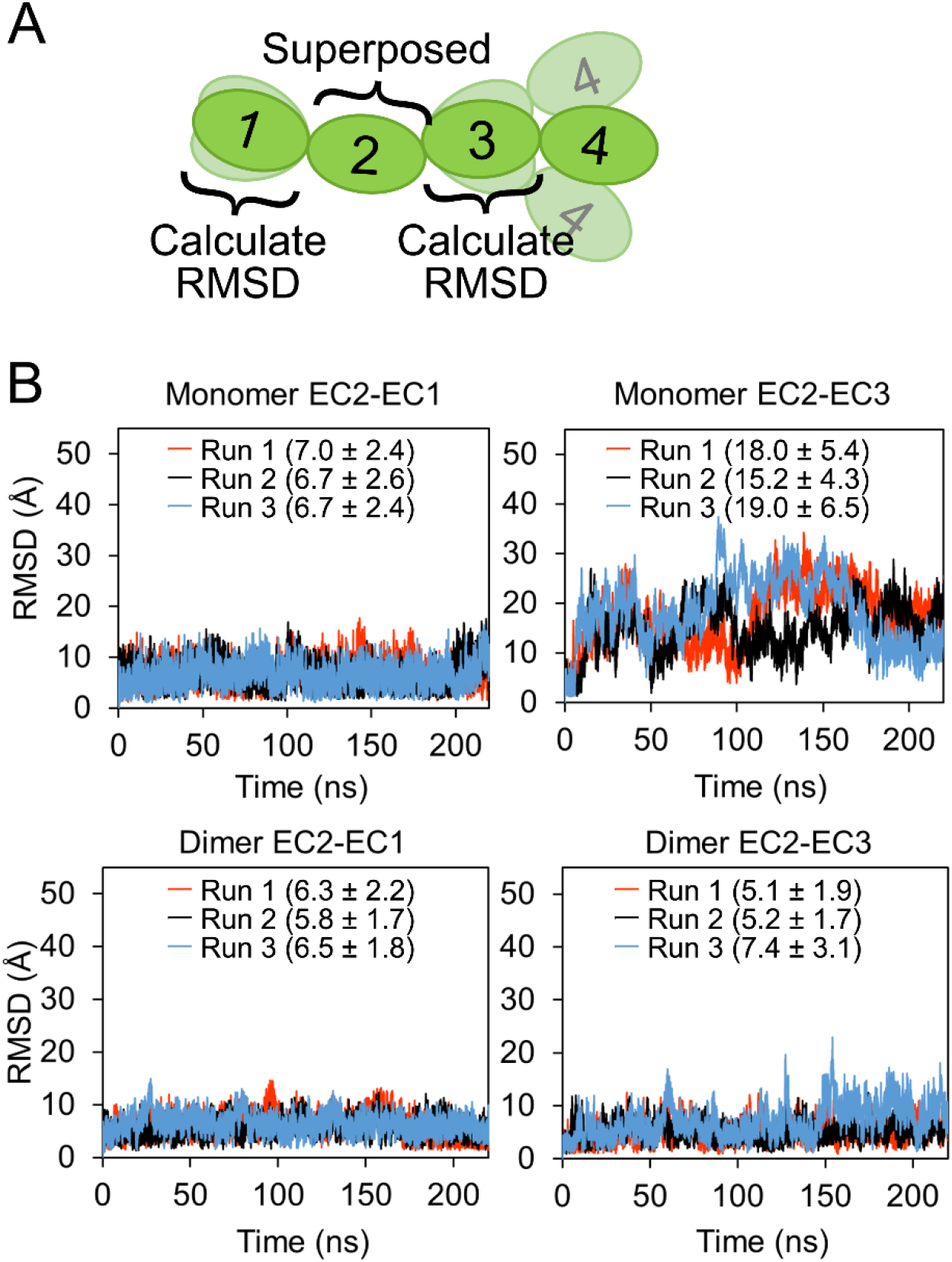
Computational analysis of the flexibility of the calcium-free linker. **A.** Schematic view of how RMSD values were calculated. **B.** RMSD values of EC1 Cα or EC3 Cα against EC2 Cα. Chain A of EC1-4 dimer structure was employed as the initial structure. **C.** RMSD values of EC1 Cα or EC3 Cα in chain A of the dimer structure against EC2 Cα in the chain A. RMSD values and standard deviations from 20 to 220 ns of each simulation are shown in parentheses in angstrom units.

Another unique characteristic in the region surrounding the calcium-free linker was the existence of an α-helix at the bottom of EC2. To the best of our knowledge, this element at the bottom of the EC2 is not found in classical cadherins. The multiple sequence alignment of the EC1-2 of human LI-, E-, N- and P-cadherin by ClustalW indicated that the insertion of the α-helix forming residues corresponded to the position immediately preceding the canonical calcium-binding motif DXE in classical cadherins (10) (Fig. S4). The Asp and Glu residues of the DXE motif in LI-cadherin dimer EC1 and EC3 coordinate with calcium ions (Fig. S5A, B) and was maintained throughout the simulation (Fig. S5C~J). The α-helix in EC2 might compensate for the absence of calcium by conferring some rigidity to the molecule.

### Interaction analysis of EC1-4 homodimer

To validate if LI-cadherin-dependent cell adhesion is mediated by the formation of homodimer observed in the crystal structure, it was necessary to find a mutant exhibiting reduced dimerization tendency. First, we analyzed the interaction between two EC1-4 molecules in the crystal structure using the PISA server (Table S5) (23). The interaction was largely mediated by EC2 of one chain of LI-cadherin and EC4 of the other chain, engaging in hydrogen bonds and non-polar contacts (Fig. 5). The dimerization surface area was 1,254 Å^2^ and a total of seven hydrogen bonds (distance between heavy atoms < 3.5 Å) were observed. Based on the analysis of these interactions, we conducted site-directed mutagenesis to assess the contribution of each residue to the dimerization of LI-cadherin. Eleven residues showing a percentage of buried area greater than 50%, or one or more intermolecular hydrogen bonds (distance between heavy atoms < 3.5 Å) were individually mutated to Ala (Table S1 and S5). To quickly identify mutants with weaker homodimerization propensity, SEC-MALS was employed.

**Figure 5.**
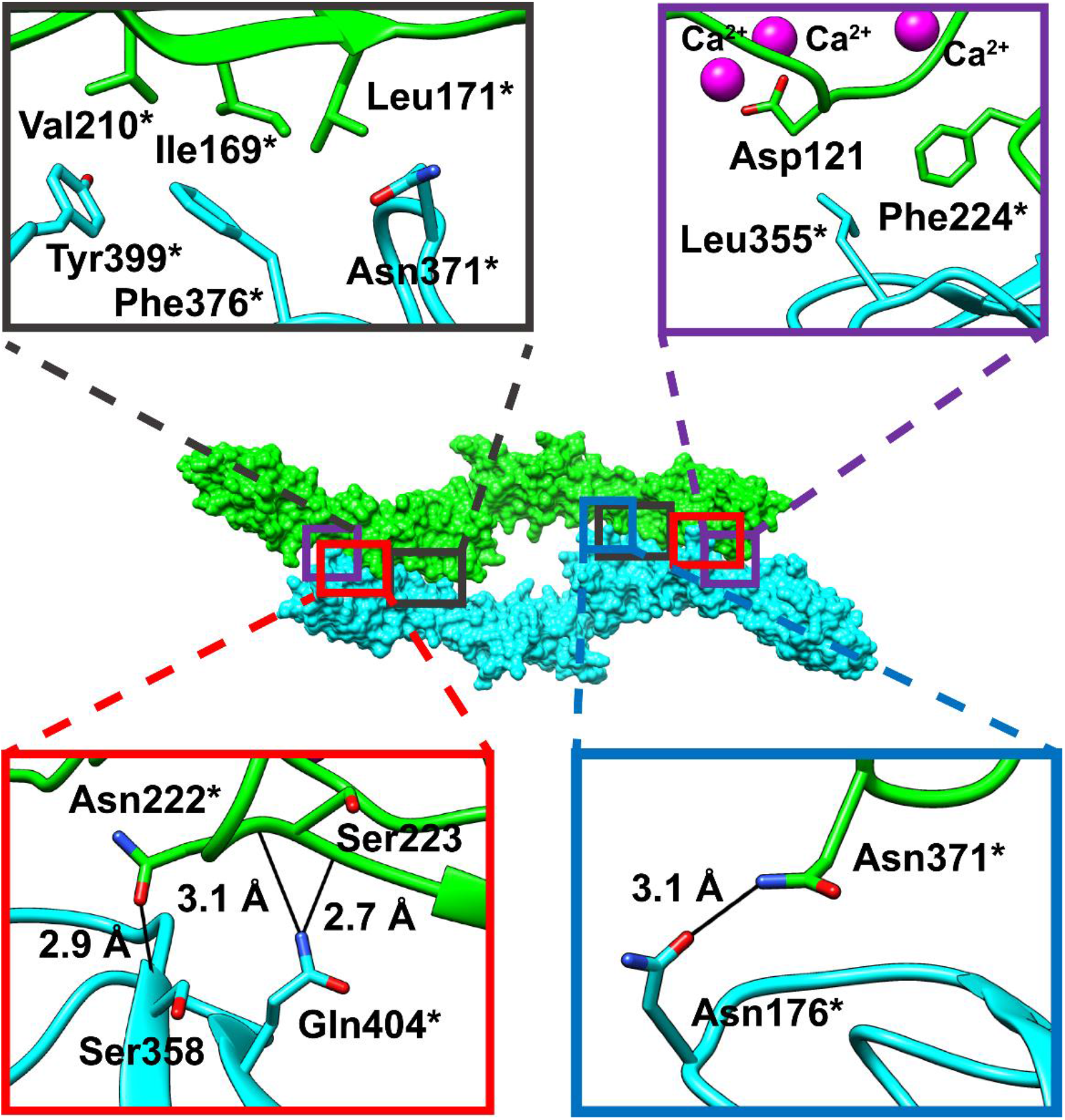
Residues involved in intermolecular interactions in the crystal structure of the EC1-4 homodimer. The non-polar interaction residues are shown in black and purple rectangles (top panels). Residues involved in hydrogen bonds (black solid lines) are shown within the red and blue rectangles (bottom panel). Residues indicated with an asterisk were individually mutated to Ala to evaluate their contribution to dimerization.

EC1-4WT (or mutants) at 100 μM were injected in the chromatographic column. Analysis of the molecular weight (MW) showed that the MW of F224A was the smallest among all the samples evaluated (Fig. 6A and Table 2). Analogous observations were made when the protein samples were injected at 50 μM (Fig. S6A). Similarly, the sample with the greatest elution volume among the 12 samples analyzed corresponded to F224A (Fig. 6B and Fig. S6B-C).

**Figure 6.**
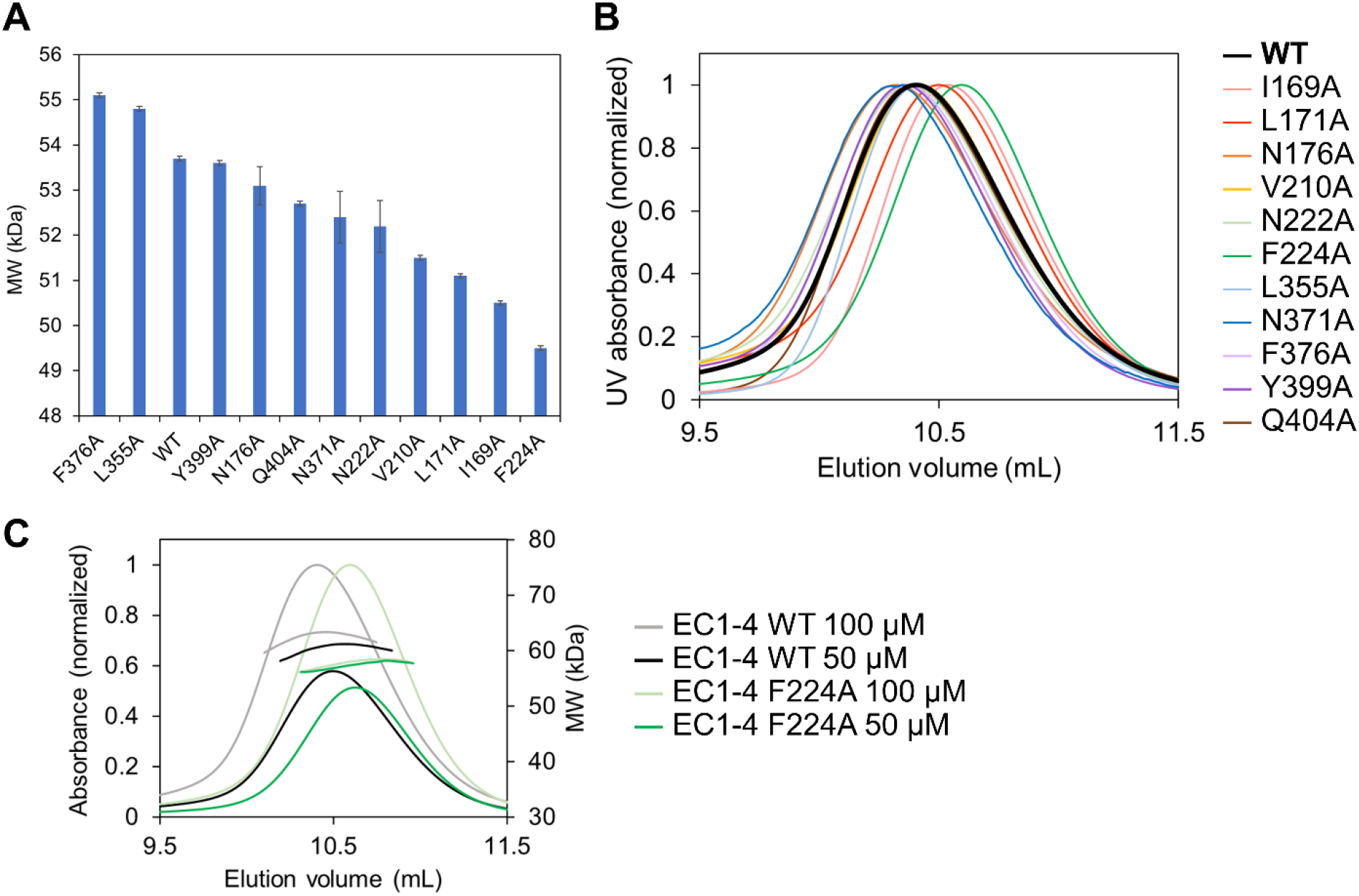
Mutagenesis analysis by SEC-MALS. **A.** Apparent molecular weight (MW) determined by MALS. The mutant F224A exhibited the smallest molecular weight among all the constructs examined. The samples were injected at 100 μM. Error bars indicate experimental uncertainties. **B.** SEC chromatograms obtained using SEC-MALS. Protein was injected at 100 μM. Chromatogram of WT and F224A are indicated in black (bold line) and green, respectively. The elution volume of F224A (determined at the peak of absorption) was the largest among all constructs inspected. **C.** SEC chromatogram and MW plots of EC1-4WT and F224A. Plots for other mutants are shown in Fig. S7C.

**Table 2:**
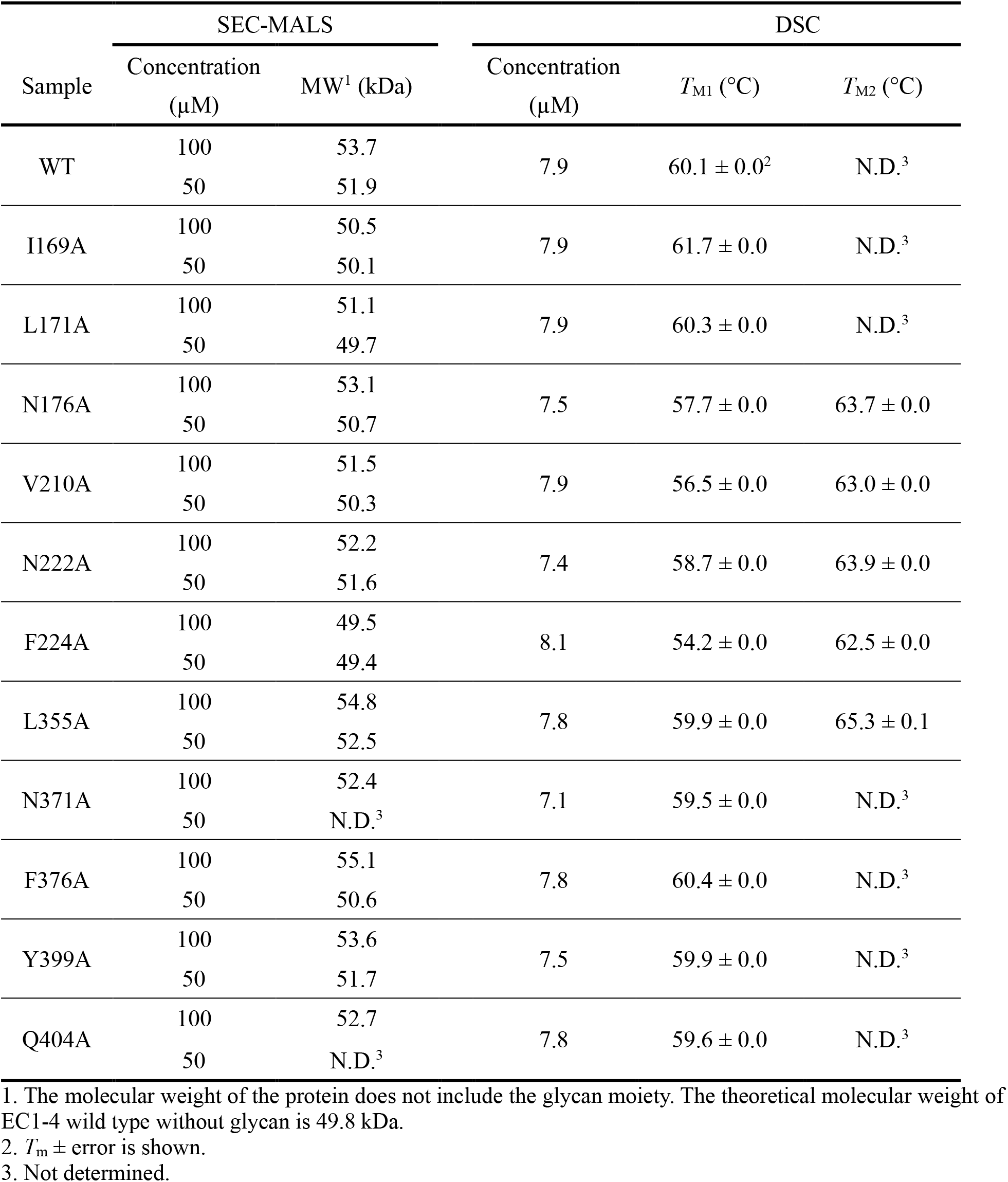
Results of Ala scanning.

We must note that the samples eluted as a single peak, corresponding to a fast equilibrium between monomers and dimers as reported in a previous study employing other cadherins (15). Although the samples were injected at 100 μM, they eluted at ~ 4 μM, since SEC will dilute the samples as they advance through the column. Considering that the *K*_D_ of dimerization of EC1-4WT determined by AUC was 39.8 μM, at a protein concentration of 4 μM, the largest fraction of the eluted sample should be monomer. This explains why the MW of the WT sample was smaller than the MW of the homodimer (99.6 kDa), and why the differences in MW among the constructs were small. We reasonably assumed that the decrease of MW and the increase of elution volume indicated a lower fraction of homodimer in the eluted sample, thus indicating a smaller dimerization tendency caused by the mutations introduced in the protein.

To confirm the disruption of the homodimer by mutation of Phe224, we performed SV-AUC measurement for EC1-4F224A. In agreement with the result of SEC-MALS, it was revealed that this construct does not form homodimer even at the highest concentration examined (120 μM, Fig. 7). Collectively, the mutational study showed that the mutation of Phe224 to Ala abolished homodimerization of EC1-4.

**Figure 7.**
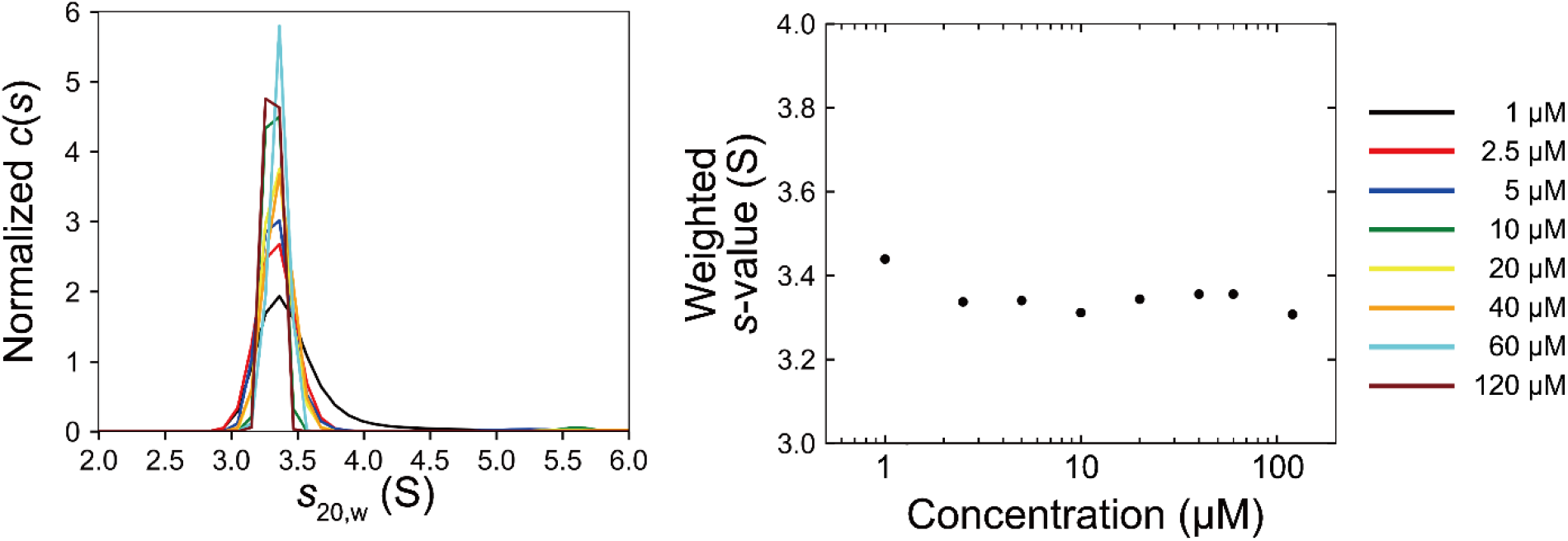
Sedimentation plot of EC1-4F224A. The signal corresponding to the dimer fraction was not observed at any of the concentrations examined. This result indicated that homodimerization of EC1-4 was abolished by the mutation of Phe224 to Ala.

### Contribution of Phe224 to dimerization

Although Phe224 does not engage in specific interactions (such as H-bonds) with the partner molecule of LI-cadherin in the crystal structure (Fig. S7A-B), it buries a significant surface (71.8 Å^2^), i.e., 95% of its total accessible surface, upon dimerization as determined by the PISA server. To understand the role of Phe224 in the dimerization of LI-cadherin, we conducted separate MD simulations of the monomeric forms of EC1-4WT and that of EC1-4F224A. We first calculated the intramolecular distance between Cα atoms of residues 224 and 122. The simulations revealed that Ala224 in the mutant moves away from the strand that contains Asn122, whereas the original Phe224 remains within a closer distance to Asn122 (Fig. 8A, Fig. S8, and Movies 3-4). The movement in the mutated residue suggests that the side chain of Phe224 engages in intramolecular interactions, being stabilized inside the pocket. Superposition of EC2 (chain A) in the crystal structure of EC1-4 and EC2 during the simulation of EC1-4F224A monomer suggests that the large movement of the loop containing Ala224 would cause steric hindrance and would inhibit dimerization (Fig. 8B).

**Figure 8.**
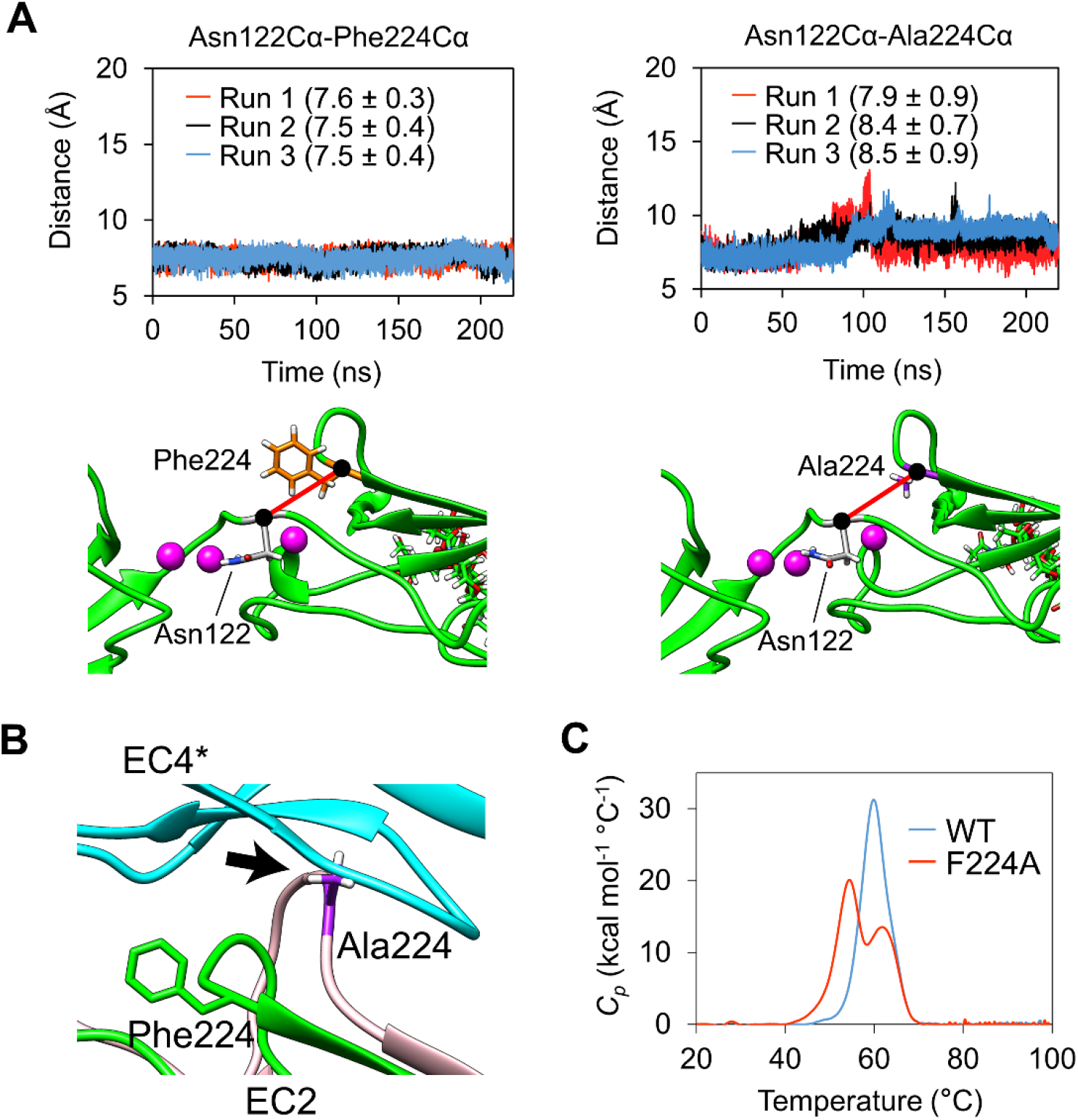
Mechanism of dimerization facilitated by Phe224. **A.** The distance between Phe224 (orange) Cα or Ala224 (purple) Cα and Asn122 (grey) Cα was evaluated by MD simulations (top panels). The Cα atoms are indicated by black circles. The distance calculated by the simulations is indicated with a thick red line joining Phe224 or Ala224 and Asn122 (bottom panels). Each MD simulation run is shown in red, black and blue. Averages and standard deviations from 20 to 220 ns of each simulation are shown in parentheses. **B.** Structure alignment of EC2 (chain A) in EC1-4 homodimer crystal structure and EC2 during the MD simulation of EC1-4F224A monomer. A snapshot of 103.61 ns in Run 1 was chosen as it showed the largest distance between Asn122 and Ala224. Ala224 is indicated in purple. The loop indicated with the black arrow would cause steric hindrance towards the formation of the homodimer. **C.** Thermal stability of EC1-4WT and F224A determined by differential scanning calorimetry. Two transitions appeared in the sample of F224A. The first transition at lower temperature seems to have appeared due to the loss of intramolecular interaction around the residue at position 224.

The analysis of the thermal stability using differential scanning calorimetry (DSC) revealed that EC1-4F224A exhibited two unfolding peaks, whereas that of EC1-4WT displayed a single peak (Fig. 8C). These results suggested that a part of EC1-4F224A molecule was destabilized by the mutation. In combination with the data from MD simulations, we propose that Phe224 contributes to the dimerization of LI-cadherin by restricting the movement of the residues around Phe224 and thus preventing the steric hindrance triggered by the large movement observed in the MD simulations of the alanine mutant. DSC measurements showed that some other mutants exhibited lower thermal stability than the wild type protein (Table 2 and Fig. S9). However, because the value of *T_M1_* of F224A is the lowest among all the mutants examined, and because other mutants displaying lower *T_M1_* than wild type did not exhibit a drastic decrease in homodimer affinity like F224A, we conclude that among the residues evaluated by Ala scanning, Phe224 was the most critical for the maintenance of homodimer structure and thermal stability.

### Functional analysis of LI-cadherin on cells

To investigate if disrupting the formation of EC1-4 homodimer influences cell adhesion, we established a CHO cell line expressing full-length LI-cadherin WT or the mutant F224A (including the transmembrane and cytoplasmic domains fused to GFP) that we termed EC1-7GFP and EC1-7F224AGFP, respectively (Table S1 and Fig. S10). We conducted cell aggregation assays and compared the cell adhesion ability of cells expressing each construct and mock cells (non-transfected Flp-In CHO) in the presence of calcium or in the presence of EDTA. The size distribution of cell aggregates was quantified using a micro-flow imaging (MFI) apparatus. EC1-7GFP showed cell aggregation ability in the presence of CaCl_2_. In contrast, EC1-7F224AGFP and mock cells did not show obvious cell aggregates in the presence of CaCl_2_ (Fig. 9A-C). From this result, it was revealed that Phe224 was crucial for LI-cadherin-dependent cell adhesion and the formation of EC1-4 homodimer in a cellular environment.

**Figure 9.**
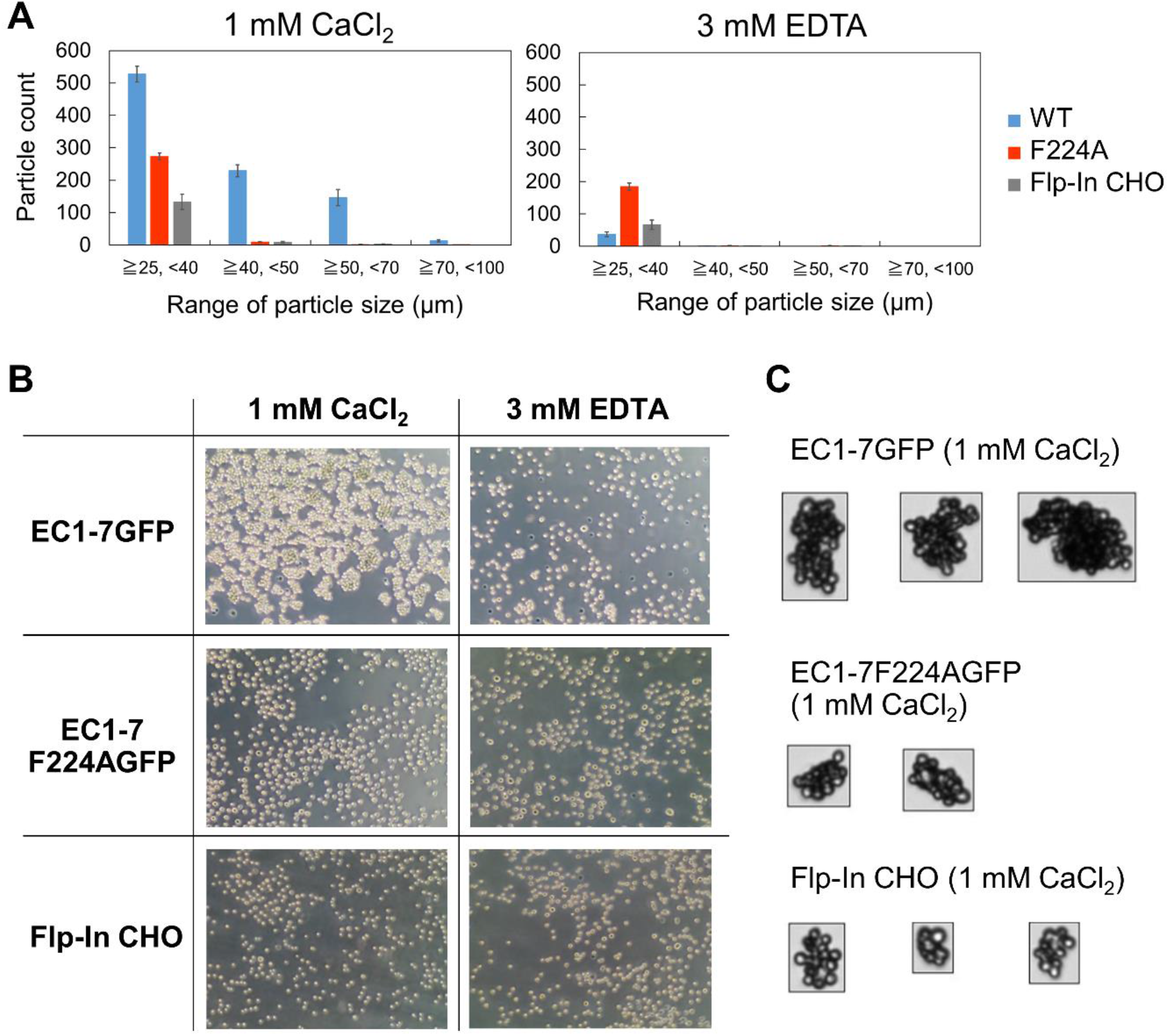
Cell aggregation assay. **A.** Size distribution of cell aggregates determined by MFI. Particles that were 25 μm or larger were regarded as cell aggregates. Only EC1-7 WT expressing cells in the existence of 1 mM CaCl_2_ showed significant number of cell aggregates that were 40 μm or larger. Data show the mean ± SEM of four measurements. **B.** Microscopy images of cell aggregates taken after adding 4% PFA and incubating the plate on ice for 20 minutes. **C.** Images of cell aggregates taken by MFI. Cell aggregates belonging to the largest size population of each construct obtained in the presence of 1 mM CaCl_2_ (70~100 μm for EC1-7GFP, 50~70 μm for EC1-7F224AGFP and 40~70 μm for Flp-In CHO) are shown.

We next performed cell aggregation assays using CHO cells expressing various constructs of LI-cadherin in which EC repeats were deleted, to elucidate the mechanism of cell-adhesion induced by LI-cadherin. LI-cadherin EC1-5 and EC3-7 expressing cells were separately established (EC1-5GFP and EC3-7GFP) (Table S1 and Fig. S10). Importantly, neither EC1-5 nor EC3-7 expressing cells showed cell aggregation ability in the presence of CaCl_2_ (Fig. 10).

**Figure 10.**
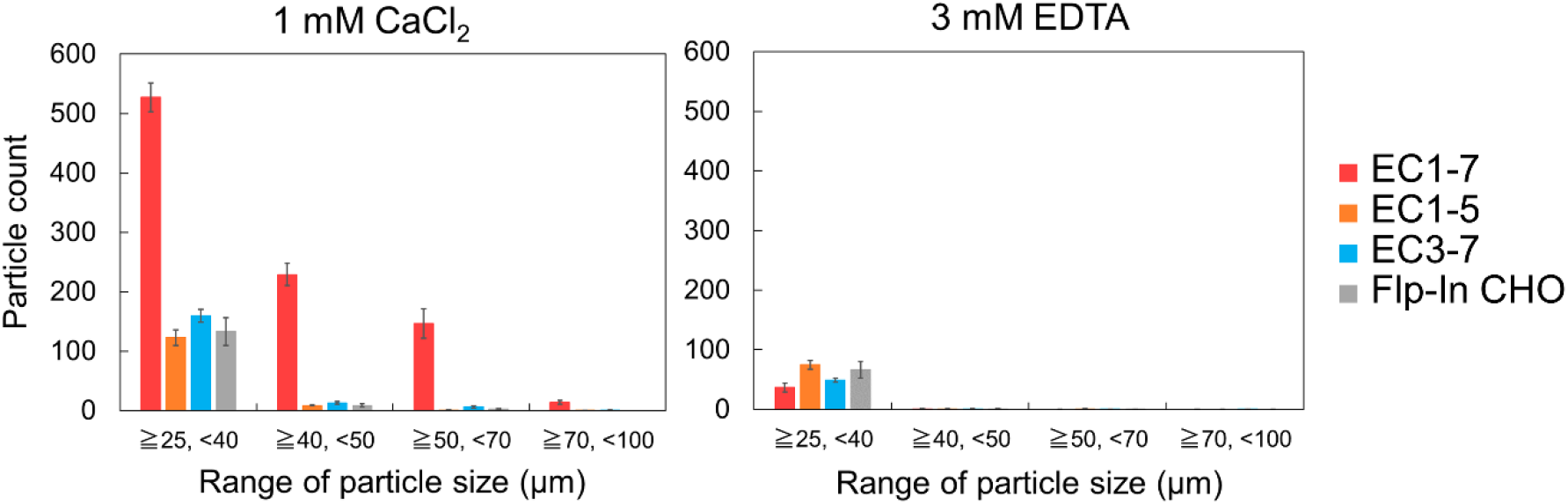
Cell adhesion mediated by short constructs. Cell aggregation assay using EC1-5GFP and EC3-7GFP. EC1-7GFP and Flp-In CHO (mock cells) were used as positive and negative control, respectively. Particles that were 25 μm or larger were considered as cell aggregates. The number of cell aggregates of both EC1-5GFP and EC3-7GFP in the presence or absence of Ca^2+^ were determined. Data are corresponded to mean ± SEM of four measurements.

Since EC1-5GFP was not conducive to cell aggregation, it is suggested that effective dimerization at the cellular level requires full-length protein. Combined with the observation from SV-AUC for EC1-7 from above, EC6 and/or EC7 should contribute to the slow dissociation of the homodimer. In the absence of EC6-7, it is likely that the dissociation rate of LI-cadherin would increase, thus impairing cell adhesion.

There are also some other alternative explanations. For example, the EC1-4 homodimer observed by X-ray crystallography and detected by AUC cannot be replicated in the EC1-5 construct in a cellular environment. We hypothesize that the overhang EC1 repeat in the dimer belonging to one cell could collide with the membrane of the opposing cell (steric hindrance) (Fig. S11A). It is also possible that inappropriate orientation of the approaching LI-cadherin molecules would contribute to the inability of EC1-5 to dimerize (Fig. S11B). An alternative possibility is that the weaker dimerization of EC1-2 (detected by SV-AUC) cannot maintain cell adhesion due to the mobility of the Ca^2+^-free linker between EC2 and EC3. Contrary to the canonical Ca^2+^-bound linker, such as the linker between EC1 and EC2, the linker between EC2 and EC3 in LI-cadherin does not contain Ca^2+^ ions. The lack of Ca^2+^ resulted in greater mobility when the EC1-4 homodimer observed by crystal structure (Fig. 3) was not formed. The combination of low dimerization affinity and high mobility likely explain the absence of EC1-2 driven cell adhesion (Fig. S11C and D).

Similarly, expression of EC3-7 on the surface of the cells did not result in cell aggregation. In this case, the observation agreed with the results of SV-AUC and SAXS, showing that EC3-4 does not dimerize. The truncation of EC1-2 from LI-cadherin generates a cadherin similar to classical cadherin in the point of view that it has five extracellular repeats and that it has a Trp residue at the N-terminus. Together with the crystal structure of the EC1-4 homodimer, which showed that Trp239 was buried in its own hydrophobic pocket and not participating in homodimerization (Fig. S12), the fact that LI-cadherin EC3-7 did not aggregate represents a unique dimerization mechanism in LI-cadherin.

EC1-5 and EC3-7 expressing cells did not show aggregation ability even when they were mixed in equal amounts (Fig. S13). This result excluded the possibility of nonsymmetrical interaction of the EC repeats (e.g., EC1-2 and EC3-4, EC1-2 and EC6-7, etc.).

## Discussion

This is the first report examining the architecture of LI-cadherin EC1-4 homodimer and the flexibility of the Ca^2+^-free linker in LI-cadherin. The mutational study and the cell aggregation assay showed that LI-cadherin-dependent cell adhesion is mediated by the formation of the dimerization interface between EC2 in one chain and EC4 in the other chain, and the contribution from other EC repeats. Our findings regarding the novel EC1-4 homodimer advances the understanding of LI-cadherin at the molecular level. The EC1-2 homodimer, which was observed by SV-AUC, and the contribution of EC6 and/or EC7 to slow the dissociation rate of the homodimer are also important to understand the mechanism of cell adhesion mediated by LI-cadherin.

The EC1-2 homodimer observed by SV-AUC appeared not to be sufficient to maintain LI-cadherin-dependent cell adhesion. Considering that there are several families of cell adhesion proteins in the human body, we cannot rule out the possibility that the EC1-2 homodimer is formed on cells where cell adhesion is maintained by other cell adhesion proteins. In LI-cadherin-dependent cell adhesion, we assume that the unique architecture of the EC1-4 homodimer was necessary to restrict the movement of the Ca^2+^-free linker and to maintain LI-cadherin-dependent cell adhesion (Fig. S14A-C).

The noncanonical α-helix in EC2 may also contribute to the unique characteristics of LI-cadherin. A previous study on the Ca^2+^-free linker of Protocadherin-15 indicated that a unique 3_10_ helix in the middle of the Ca^2+^-free linker is one of the factors conferring mechanical strength to the linker (21). We assume that the α-helix in LI-cadherin EC2 is required to maintain structural rigidity in the absence of the coordination of negatively charged residues to Ca^2+^.

Contribution of EC6 and/or EC7 on homodimerization was also a notable factor. C-cadherin is known to form strand-swap dimer which is mediated by interactions at the N-terminal strand of EC1 (18). However, bead aggregation assay and laminar flow assay suggested that C-cadherin-dependent binding activity is maintained through interactions of multiple EC repeats (24). Likewise, our findings suggested that dimerization of LI-cadherin results from a collective effort from several repeats throughout the protein. According to data from the cell aggregation assays of the mutant EC1-7F224AGFP, it is clear that LI-cadherin-dependent cell adhesion is abolished by the mutation of Phe224, a residue located within the EC2 repeat. This result suggests that EC6 and/or EC7 contribute to cell adhesion after the formation of the EC1-4 homodimer, an idea that is also supported by the observation that EC3-7GFP does not show cell aggregation ability. The mechanism underlying the contribution of EC6 and EC7 repeats to the slow dissociation rate of the EC1-7 homodimer needs to be further investigated.

Several differences between LI-cadherin and E-cadherin might explain the reason for the unique biological characteristics of LI-cadherin. Both LI-cadherin and E-cadherin are expressed on normal intestine cells, however, their sites of expression are different. LI-cadherin is expressed at the intercellular cleft and is excluded from the adherens junctions (7), where E-cadherin is precisely expressed (25). Even though LI-cadherin is not present at the adherens junctions, trans-interaction of LI-cadherin is necessary to maintain water transport through the intercellular cleft of intestine cells (8). Clustering on cell membrane might also be different. Classical cadherins including E-cadherin are thought to form clusters on the cell membrane to facilitate cell adhesion (18). The lateral interaction interface of these cadherins was suggested from packing contacts in the lattice of protein crystals. In contrast to classical cadherins, we did not observe crystal packing contacts that might suggest lateral (cis) interactions in our crystal structure. Indeed, our crystal structure indicates that the few N-glycans present in LI-cadherin are directed towards the opposite side of the homodimer interface, suggesting that the protein chains belonging to the homodimer do not participate in cis-interactions. Considering that the interface area of the X-dimer and the strand-swap dimer are smaller than that of LI-cadherin EC1-4 dimer, we speculate that LI-cadherin form homodimers with a broader interface which could maintain trans-interactions without the need of cis-clusters on the cell membrane.

Expression of LI-cadherin is observed on various cancer cells such as gastric adenocarcinoma, colorectal cancer cells and pancreatic cancer cells (4, 26, 27). The roles of LI-cadherin on cancer cells have been discussed previously. For example, it was shown that inoculation of LI-cadherin gene (CDH17)-silenced cells in nude mice inhibited the progression of colorectal cancer (28). In case of gastric cancer, the size of LI-cadherin-positive tumor was significantly larger than that of LI-cadherin-negative tumor (29). Considering that loss of cell adhesion ability by the downregulation of E-cadherin by epithelial mesenchymal transition (EMT) is often observed in cancer cells (30, 31), the fact that LI-cadherin is upregulated in various types of cancer cells suggest that LI-cadherin acts differently with E-cadherin on cancer cells. The unique architecture of the LI-cadherin homodimer and the absence of interactions with intracellular (cytoplasmic) proteins (12) suggest a distinctive role of LI-cadherin in cancer cells with respect to that of classical cadherins.

In summary, our study with LI-cadherin has unveiled novel molecular-level features for the dimerization of a cadherin molecule. The data suggest that molecules targeting interface of LI-cadherin homodimer abrogate the LI-cadherin-dependent cell adhesion. On the other hand, we estimate that molecules that restrict the movement of Ca^2+^-free linker might strengthen LI-cadherin-dependent cell adhesion by stabilizing the LI-cadherin homodimer.

## Experimental procedures

### Protein sequence

Amino acid sequence of recombinant protein and LI-cadherin expressing CHO cells are summarized in Table S1.

### Expression and purification of recombinant LI-cadherin

All LI-cadherin constructs were expressed using the same method. All constructs were cloned in pcDNA 3.4 vector (ThermoFisher Scientific). Recombinant protein was expressed using Expi293F™ Cells (ThermoFisher Scientific) following manufacturer’s protocol. Cells were cultured for three days after transfection at 37 °C and 8% CO_2_.

Purification method was identical for all the constructs except for EC1-7 (see below). The supernatant was collected and filtered followed by dialysis against a solution composed of 20 mM Tris-HCl at pH 8.0, 300 mM NaCl, and 3 mM CaCl_2_. Immobilized metal affinity chromatography (IMAC) was performed using Ni-NTA Agarose (Qiagen). Protein was eluted by 20 mM Tris-HCl at pH 8.0, 300 mM NaCl, 3 mM CaCl_2_, and 300 mM Imidazole. Final purification was performed by size exclusion chromatography (SEC) using HiLoad 26/600 Superdex 200 pg column (Cytiva) at 4 °C equilibrated in buffer A (10 mM HEPES-NaOH at pH 7.5, 150 mM NaCl, and 3 mM CaCl_2_). Unless otherwise specified, samples were dialyzed against buffer A before analysis and filtered dialysis buffer was used for assays.

For the purification of EC1-7, dialysis after the collection of the supernatant and IMAC were performed by the same method as other constructs. After the purification with IMAC, the fraction containing protein was dialyzed against 20 mM Tris-HCl, pH 8.0, 5 mM NaCl, and 3 mM CaCl_2_, and anion exchange chromatography was performed using a HiTrap Q HP column (1-mL size; Cytiva). The column was washed with Anion A buffer (20 mM Tris-HCl, pH 8.0, 10 mM NaCl, and 3 mM CaCl_2_) prior to the injection of the protein. The percentage of Anion B buffer (20 mM Tris-HCl, pH 8.0, 500 mM NaCl, and 3 mM CaCl_2_) was increased in the stepwise manner in increments of 12.5% to elute the protein. Elution at an anion B buffer percentage of approximately 37.5% to 50% was collected for final purification. The final purification was performed by injecting the collected fractions onto a HiLoad 26/600 Superdex 200 pg column (Cytiva) at 4 °C equilibrated in buffer A.

### Sedimentation velocity analytical ultracentrifugation (SV-AUC)

SV-AUC experiments were conducted using the Optima AUC (Beckman Coulter) equipped with an 8-hole An50 Ti rotor at 20 °C with LI-cadherin constructs dissolved in buffer A. Measurements of EC1-7 were performed at 1, 2.5, 5, 10, and 20 μM. Measurements of EC1-2, EC3-4, EC1-4, and EC1-5 were performed at 1, 2.5, 5, 10, 20, 40, and 60 μM. Measurements of EC1-4F224A were performed at 1, 2.5, 5, 10, 20, 40, 60, and 120 μM. Protein sample (390 μL) was loaded into the sample sector of a cell equipped with sapphire windows and 12 mm double-sector charcoal-filled upon centerpiece. A volume of 400 μL of buffer was loaded into the reference sector of each cell. Data were collected at 42,000 rpm with a radial increment of 10 μm using a UV detection system.

The collected data were analyzed using continuous *c*(*s*) distribution model implemented in program SEDFIT (version 16.2b) (32) fitting for the frictional ratio, meniscus, time-invariant noise, and radial-invariant noise using a regularization level of 0.68. The sedimentation coefficient ranges of 0-15 S were evaluated with a resolution of 150. The partial specific volumes of EC1-7, EC1-2, EC3-4, EC1-4, EC1-5, and EC1-4F224A were calculated based on the amino acid composition of each sample using program SEDNTERP 1.09 (33) and were 0.732 cm^3^/g, 0.730 cm^3^/g, 0.733 cm^3^/g, 0.732 cm^3^/g, and 0.734 cm^3^/g, and 0.732 cm^3^/g, respectively. The buffer density and viscosity were calculated using program SEDNTERP 1.09 as 1.0055 g/cm^3^ and 1.025 cP, respectively. Figures of *c*(*s_20, w_*) distribution were generated using program GUSSI (version 1.3.2) (34). The weight-average sedimentation coefficient of each sample was calculated by integrating the range of sedimentation coefficients where peaks with obvious concentration dependence were observed. For the determination of the dissociation constant of monomer-dimer equilibrium, *K*_D_, the concentration dependence of the weight-average sedimentation coefficient was fitted to the monomer-dimer self-association model implemented in program SEDPHAT (version 15.2b) (35).

### Solution structure analysis using SAXS

All measurements were performed at beamline BL-10C (36) of the Photon Factory (Tsukuba, Japan). The experimental procedure is described previously (19). Concentrations of EC3-4 was 157 μM. Data were collected using a PILATUS3 2M (Dectris) and image data were processed by SAngler software (37). A wavelength was 1.488 Å with a camera distance 101 cm. Exposure time was 60 seconds and raw data between *s* values of 0.010 and 0.84 Å^-1^ were measured. The background scattering intensity of buffer was subtracted from each measurement. The scattering intensities of four measurements were averaged to produce the scattering curve of EC3-4. Data are placed on an absolute intensity scale. Conversion factor was calculated based on the scattering intensity of water. The calculation of the theoretical curves of SAXS and χ^2^ values were performed using FoXS server (38, 39).

### MD simulations

Molecular dynamics simulations of LI-cadherin were performed using GROMACS 2016.3 (40) with the CHARMM36m force field (41). A whole crystal structure of EC1-4 homodimer, EC1-4 monomer form, EC1-4F224A monomer form and EC3-4 monomer form was used as the initial structure of the simulations, respectively. EC1-4 and EC3-4 of chain A was extracted from EC1-4 homodimer crystal structure to generate EC1-4 monomer form and EC3-4 monomer form, respectively. Sugar chains were removed from the original crystal structure. Missing residues were modelled by MODELLER 9.18 (42). Solvation of the structures were performed with TIP3P water (43) in a rectangular box such that the minimum distance to the edge of the box was 15 Å under periodic boundary conditions through the CHARMM-GUI (44). Addition of N-bound type sugar chains (G0F) and the mutation of Phe224 in EC1-4 monomer to Ala224 were also performed through the CHARMM-GUI (44, 45). The protein charge was neutralized with added Na or Cl, and additional ions were added to imitate a salt solution of concentration 0.15 M. Each system was energy-minimized for 5000 steps and equilibrated with the NVT ensemble (298 K) for 1 ns. Further simulations were performed with the NPT ensemble at 298 K. The time step was set to 2 fs throughout the simulations. A cutoff distance of 12 Å was used for Coulomb and van der Waals interactions. Long-range electrostatic interactions were evaluated by means of the particle mesh Ewald method (46). Covalent bonds involving hydrogen atoms were constrained by the LINCS algorithm (47). A snapshot was saved every 10 ps. All trajectories were analyzed using GROMACS tools. RMSD, dihedral angles, distances between two atoms and clustering were computed by rms, gangle, distance and cluster modules, respectively.

The convergence of the trajectories was confirmed by calculating RMSD values of Cα atoms (Fig. S4A, B and S7A, B). As the molecule showed high flexibility at Ca^2+^-free linker, as for EC1-4WT monomer, EC1-4F224A monomer and EC1-4 dimer, RMSD of each EC repeat was calculated individually. Five Cα atoms at N-terminus were excluded from the calculation of RMSD of EC1 as they were disordered. As the RMSD values were stable after running 20 ns of simulations, we did not consider the first 20 ns when we analyzed the trajectories.

### Generation of EC3-4_plus

MD simulation of the EC3-4 monomer was performed for 220 ns. The trajectories from 20 ns to 220 ns were clustered using the ‘cluster’ tool of GROMACS. The structure which exhibited the smallest average RMSD from all other structures of the largest cluster was termed EC3-4_plus and used for the purpose of comparison with the data in solution (SAXS).

### Crystallization of LI-cadherin EC1-4

Purified LI-cadherin EC1-4 was dialyzed against 10 mM HEPES-NaOH at pH 7.5, 30 mM NaCl, and 3 mM CaCl_2_ prior to crystallization. After the dialysis, the protein was concentrated to 314 μM. Optimal condition for crystallization was screened using an Oryx8 instrument (Douglas Instruments) using commercial screening kits (Hampton Research). The crystal used for data collection was obtained in a crystallization solution containing 200 mM sodium sulfate decahydrate and 20% w/v Polyethylene glycol 3,350 at 20 °C. Suitable crystals were harvested, briefly incubated in mother liquor supplemented with 20% glycerol, and transferred to liquid nitrogen for storage until data collection.

### Data collection and refinement

Diffraction data from single crystals of EC1-4 were collected in beamline BL-5A at the Photon Factory (Tsukuba, Japan) under cryogenic conditions (100 K). Diffraction images were processed with the program MOSFLM and merged and scaled with the program SCALA (48) of the CCP4 suite (49). The structure of EC1-4 was determined by the method of molecular replacement using the coordinates of P-cadherin (PDB entry code 4ZMY) (50) and LI-cadherin EC1-2 (PDB entry code 7EV1) with the program PHASER (51). The model thus obtained was further refined with the programs REFMAC5 (52) and extensively built with COOT (53). Validation was carried out with PROCHECK (54). Data collection and structure refinement statistics are given in Table 1. UCSF Chimera was used to render all of the molecular graphics (55).

### Site-directed mutagenesis

Site-directed mutagenesis was performed as described previously (56).

### Size exclusion chromatography with multi-angle light scattering (SEC-MALS)

The molecular weight of LI-cadherin was determined using Superose 12 10/300 GL column (Cytiva) with inline DAWN8+ multi angle light scattering (MALS) (Wyatt Technology), UV detector (Shimadzu), and refractive index (RI) detector (Shodex). Protein samples (45 μL) were injected at 100 μM or 50 μM. Analysis was performed using ASTRA software (Wyatt Technology). Concentration at the end of the chromatographic column was measured based on the UV absorbance. The protein conjugate method was employed for the analysis as sugar chains were bound to LI-cadherin. All detectors were calibrated using bovine serum albumin (BSA) (Sigma-Aldrich).

### Comparison of thermal stability by DSC

DSC measurement was performed using MicroCal VP-Capillary DSC (Malvern). The measurement was performed from 10 °C to 100 °C at the scan rate of 1 °C min^-1^. Data was analyzed using Origin7 software.

### Establishment of CHO cells expressing LI-cadherin

The DNA sequence of monomeric GFP was fused at the C-terminal of all human LI-cadherin constructs of which stable cell lines were established and was cloned in pcDNA™5/FRT vector (ThermoFisher Scientific). CHO cells stably expressing LI-cadherin-GFP were established using Flp-In™-CHO Cell Line following the manufacturer’s protocol (ThermoFisher Scientific). Cloning was performed by the limiting dilution-culture method. Cells expressing GFP were selected and cultivated. Observation of the cells were performed by In Cell Analyzer 2000 (Cytiva). The cells were cultivated in Ham’s F-12 Nutrient Mixture (ThermoFisher Scientific) supplemented with 10 % fetal bovine serum (FBS), 1% L-Glutamine or 1% GlutaMAX™-I (ThermoFisher Scientific), 1% penicillin-streptomycin, and 0.5 mg mL^-1^ Hygromycin B at 37 °C and 5.0% CO_2_.

### Cell Imaging

Cells (100 μL) were added to a 96-well plate (Greiner) at 1 x 10^5^ cells mL^-1^ and cultured overnight. After washing the cells with wash medium (Ham’s F-12 Nutrient Mixture (ThermoFisher Scientific) supplemented with 10 % fetal bovine serum (FBS), 1% GlutaMAX™-I, 1%penicillin-streptomycin), Hoechst 33342 (ThermoFisher Scientific) (100 μL) was added to each well at 0.25 μg mL^-1^. The plate was incubated at room temperature for 30 minutes. Cells were washed with wash medium twice and with 1x HMF (10 mM HEPES-NaOH at pH 7.5, 137 mM NaCl, 5.4 mM KCl, 0.34 mM Na_2_HPO_4_, 1 mM CaCl_2_, and 5.5 mM glucose) twice. After that, 1x HMF (200 μL) was loaded to each well and the images were taken with an In Cell Analyzer 2000 instrument (Cytiva) using the FITC filter (490/20 excitation, 525/36 emission) and the DAPI filter (350/50 excitation, 455/50 emission) with 60 x 0.70 NA objective lens (Nikon).

### Cell aggregation assay

Cell aggregation assay was performed by modifying the methods described previously (57, 58). Cells were detached from cell culture plate by adding 1x HMF supplemented with 0.01% trypsin and placing on a shaker at 80 rpm for 15 minutes at 37 °C. FBS was added to the final concentration of 20% to stop the trypsinization. Cells were washed with 1x HMF supplemented with 20%FBS once and with 1x HMF twice to remove trypsin. Cells were suspended in 1x HMF at 1 x 10^5^ cells mL^-1^. 500 μL of the cell suspension was loaded into 24-well plate coated with 1% w/v BSA. EDTA was added if necessary. After incubating the plate at room temperature for 5 minutes, 24-well plate was placed on a shaker at 80 rpm for 60 minutes at 37 °C.

### Micro-Flow Imaging (MFI)

Micro-Flow Imaging (Brightwell Technologies) was used to count the particle number and to visualize the cell aggregates after cell aggregation assay. After the cell aggregation assay described above, the plate was incubated at room temperature for 10 minutes and 500 μL of 4% Paraformaldehyde Phosphate Buffer Solution (Nacalai Tesque) was loaded to each well. The plate was incubated on ice for more than 20 minutes. Images of the cells were taken using EVOS® XL Core Imaging System (ThermoFisher Scientific) if necessary. After that, cells were injected to MFI.

MFI View System Software was used for the measurements and analyses. The instrument was flushed with detergent and ultrapure water prior to the experiments. Cleanliness of the flow channel was checked by performing the measurement using ultrapure water and confirming that less than 100 particles/mL was detected. Flow path was washed with 1x HMF before the measurements of the samples. Purge volume and analyzed volume were 200 μL and 420 μL, respectively. Optimize Illumination was performed prior to each measurement. Particles within a size of 100 to 1 μm were counted.

### Data availability

The coordinates and structure factors of LI-Cadherin EC1-4 have been deposited in the Protein Data Bank with entry code 7CYM. All remaining data are contained within the article.

## Acknowledgements

We thank Dr. S. Kudo and Dr. H. Akiba for expert advice. We thank Dr. O. Kusano-Arai, Dr. H. Iwanari and Dr. T. Hamakubo for providing us with gene sequence of LI-cadherin.

## Funding and additional information

The supercomputing resources in this study were provided by the Human Genome Center at the Institute of Medical Science, The University of Tokyo, Japan. This work was funded by a Grant-in-Aid for Scientific Research (A) 16H02420 (K.T.) and a Grant-in-Aid for Scientific Research (B) 20H02531 (K.T.) from Japan Society for the Promotion of Science, a Grant-in-Aid for Scientific Research on Innovative Areas 19H05760 and 19H05766 (K.T.) from Ministry of Education, Culture, Sports, Science and Technology, and a Grant-in-Aid for JSPS fellows 18J22330 (A.Y.) from Japan Society for the Promotion of Science. We are grateful to the staff of the Photon Factory (Tsukuba, Japan) for excellent technical support. Access to beamlines BL-5A and BL-10C was granted by the Photon Factory Advisory Committee (Proposal Numbers 2018G116 and 2017G661).

