## Supporting Information for "Dimerization mechanism and structural features of human LI-cadherin"

**Running title**: Dimerization mechanism of LI-cadherin

**Table S1: Composition of the constructs used in this study.**

**Table S2: Sequence homology between human LI-cadherin EC3-7 (residues 238-777)[a] and human E-, N-, and P-cadherin.**

**Table S3: Sequence homology between human LI-cadherin EC1-2 (residues 23-237)[a] and human E-, N-, and P-cadherin.**

**Table S4: Parameters and data of the SAXS measurement of EC3-4.**

**Table S5: Interaction surface in LI-cadherin homodimer.**

**Figure S1. SAXS data of EC3-4.**

**Figure S2. RMSD of Cα atoms during the simulations of EC1-4 WT monomer and homodimer.**

**Figure S3. Dihedral angles of each domain.**

**Figure S4. Sequence alignment of EC1-2 of human LI-, E-, N- and P-cadherin.**

**Figure S5. Coordination of Asp and Glu residues to Calcium ions.**

**Figure S6. Results of SEC-MALS when the samples were injected at 50 μM.**

**Figure S7. Residues interacting with Phe224 in chain A.**

**Figure S8. RMSD of Cα atoms during the simulations of EC1-4F224A monomer.**

**Figure S9. DSC data corresponding to EC1-4 mutants.**

**Figure S10. Verification of expression of LI-cadherin in mammalian cells.**

**Figure S11. Schematic views of LI-cadherin EC1-5 homodimer.**

**Figure S12. Residues interacting with Trp239 in chain A depicted by LigPlot+.**

**Figure S13. Size distribution of cell aggregates measured by MFI.**

**Figure S14. Schematic view of homodimer formed by LI-cadherin.**

**Movie 1. A simulation of EC1-4 monomer.**

**Movie 2. A simulation of EC1-4 dimer.**

**Movie 3. Magnified video image of Asn122 and Phe224 during a simulation of EC1-4 WT monomer.**

**Movie 4. Magnified video image of Asn122 and Ala224 during a simulation of EC1-4 F224A monomer.**

**Table S1: Composition of the constructs used in this study.**

| Construct Name | Sequence[c] | Mutation |
|---|---|---|
| EC1-7[a] | 23-777 | × |
| EC1-5[a] | 23-563 | × |
| EC1-4[a] | 23-441 | × |
| EC1-4I169A[a] | | I169A |
| EC1-4L171A[a] | | L171A |
| EC1-4N176A[a] | | N176A |
| EC1-4V210A[a] | | V210A |
| EC1-4N222A[a] | | N222A |
| EC1-4F224A[a] | | F224A |
| EC1-4L355A[a] | | L355A |
| EC1-4N371A[a] | | N371A |
| EC1-4F376A[a] | | F376A |
| EC1-4Y399A[a] | | Y399A |
| EC1-4Q404A[a] | | Q404A |
| EC1-2[a] | 23-236 | × |
| EC3-4 (AUC)[a] | 238-441 | × |
| EC3-4 (SAXS)[a] | 238-449 | × |
| EC1-7GFP[b] | 23-832 | × |
| EC1-7F224AGFP[b] | | F224A |
| EC1-5GFP[b] | 23-558, 778-832 | × |
| EC3-7GFP[b] | 238-832 | × |

a: Myc-tag (EQKLISEEDL) followed with NSAVD sequence and His-tag (HHHHHH) were added to the C-terminus.
b: Gly-Ser linker and monomeric GFP were added to the C-terminus.
c: Amino acid numbering follows the numbering of Uniprot entry Q12864.

**Table S2: Sequence homology between human LI-cadherin EC3-7 (residues 238-777)[a] and human E-, N-, and P-cadherin.**

| Construct | Sequence[a] | Homology (%) |
|---|---|---|
| E-cadherin EC1-5 | 155-709 | 29.9 |
| N-cadherin EC1-5 | 160-724 | 30.1 |
| P-cadherin EC1-5 | 108-654 | 28.3 |

a: Amino acid numbering of LI-, E-, N-, and P-cadherin follows the numbering of Uniprot entry Q12864, P12830, P19022, and P22223, respectively.

**Table S3: Sequence homology between human LI-cadherin EC1-2 (residues 23-237)[a] and human E-, N-, and P-cadherin.**

| Construct | Sequence[a] | Homology (%) |
|---|---|---|
| E-cadherin EC1-2 | 155-371 | 31.2 |
| N-cadherin EC1-2 | 160-378 | 32.0 |
| P-cadherin EC1-2 | 108-324 | 31.8 |

a: Amino acid numbering of LI-, E-, N-, and P-cadherin follows the numbering of Uniprot entry Q12864, P12830, P19022, and P22223, respectively.

**Table S4: Parameters and data of the SAXS measurement of EC3-4.**

| **Experimental parameters** | |
|---|---|
| Concentration (µM) | 157 |
| Exposure time (sec) | 60 |
| **Experimental data** | |
| *I*(0) (absolute scale) ($cm^{-1}$)[a] | 0.30 |
| *Rg* (Å) (Guinier)[b] | 32.97 |
| $D_{max}$ (Å)[b] | 127 |
| MW(Qp) (kDa)[c] | 29.7 |
| MW(MoW) (kDa)[c] | 33.6 |
| MW(Vc) (kDa)[c] | 28.3 |

a: Water was used as a calibrant to convert scattering intensity into absolute intensity.

b: Guinier plot and pair distance distribution function (p(r) function) are shown in Fig. S1. Guinier plot and p(r) function were generated using AUTORG (59) and GNOM (60), respectively.

c: MW was calculated by DATMW (61–63).

**Table S5: Interaction surface in LI-cadherin homodimer.**

| Residue[a] | ASA[b] (Å$^2$) | BSA[c] (Å$^2$) | BAP[d] (%) | Hydrogen bond, atom involving in H-bond | Distance[e] (Å) | Hydrogen bond partner[f] |
|---|---|---|---|---|---|---|
| Ile169 | 72.32 | 43.37 | 59.97 | × | — | — |
| Leu171 | 94.84 | 74.72 | 78.79 | × | — | — |
| Asn176[g] | 128.81 | 15.43 | 11.98 | × | — | — |
| Val210 | 37.89 | 35.20 | 92.90 | × | — | — |
| Asn222 | 125.44 | 63.76 | 50.83 | ○, OD1 | 3.59 | Asn357-ND2 |
| | | | | ○, OD1 | 2.88 | Ser358-N |
| | | | | ○, O | 3.06 | Gln404-NE2 |
| Phe224 | 74.55 | 70.33 | 94.34 | × | — | — |
| Leu355 | 115.89 | 69.43 | 59.91 | × | — | — |
| Asn371 | 156.99 | 91.14 | 58.05 | ○, ND2 | 3.11 | Asn176-OD1 |
| Phe376 | 121.18 | 75.20 | 62.06 | × | — | — |
| Tyr399 | 133.27 | 71.17 | 53.40 | × | — | — |
| Gln404 | 42.59 | 41.48 | 97.39 | ○, NE2 | 3.18 | Asn222-O |
| | | | | ○, NE2 | 2.77 | Ser223-O |

a: All residues belong to chain A.

b: ASA: Accessible Surface Area

c: BSA: Buried Surface Area

d: BAP: Buried Area Percentage

e: Distance between heavy atoms involved in H-bond formation.

f: All residues belong to chain B.

g: As Asn176 side chain formed hydrogen bond with the residue in chain B, mutation analysis of Asn176 was performed even though BAP is lower than 50%.

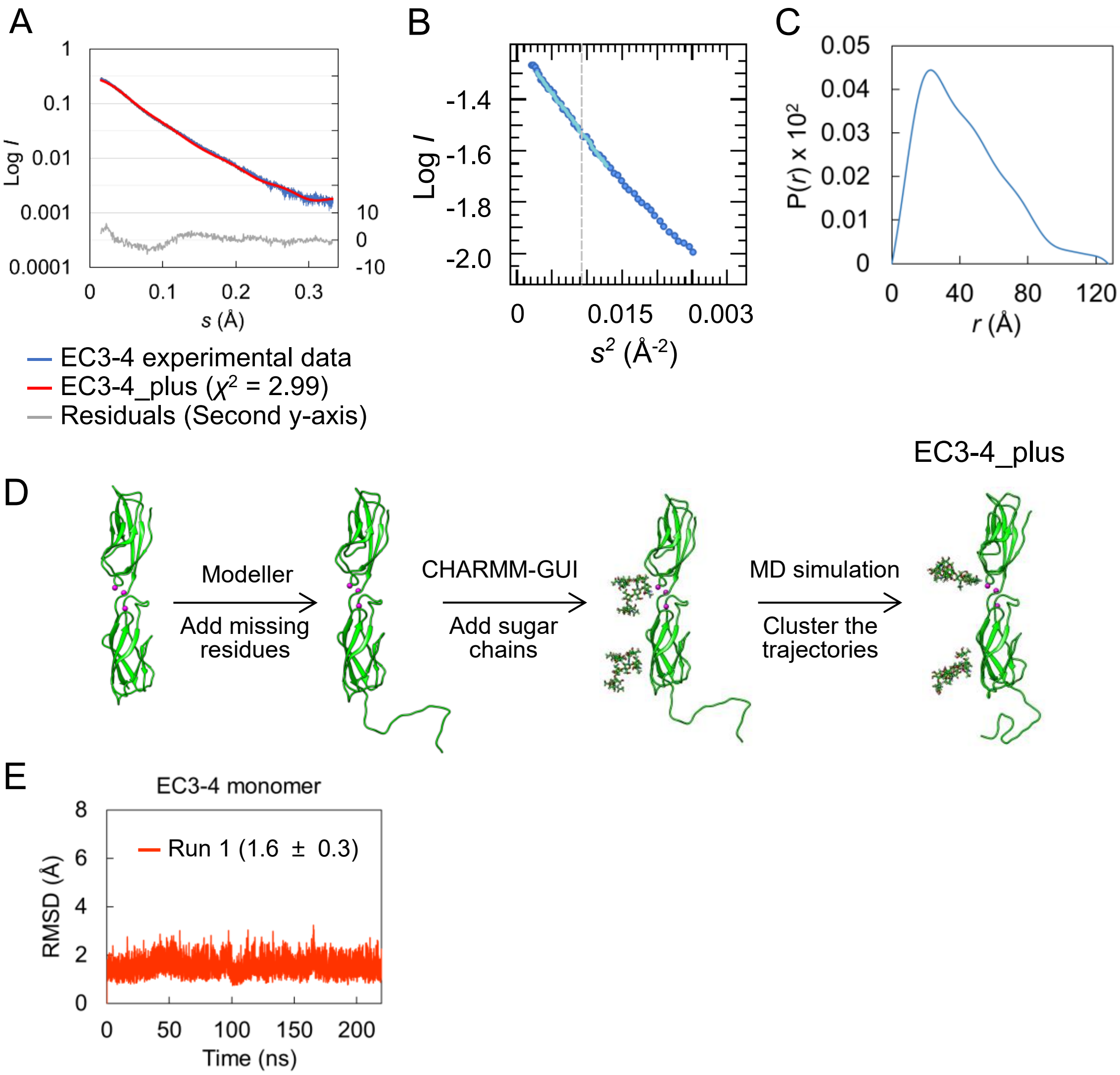


**Figure S1.** SAXS data of EC3-4. **A.** Experimental and theoretical curves along with the residuals. The theoretical curve was calculated using the FoXS server. The method to generate the modified crystal structure is described in experimental procedures and supplementary information. **B.** Guinier plot of EC3-4. Experimental data and fitting are indicated in blue and light blue, respectively. **C.** P(r) function of EC3-4. **D.** Procedures to generate EC3-4_plus. Sugar chains were removed from the original crystal structure. Missing residues and sugar chain were modelled using MODELLER 9.18 and CHARMM-GUI, respectively. MD simulation was performed for 220 ns and the trajectories from 20 ns to 220 ns were clustered using GROMACS tool. The center structure of the largest cluster was determined as EC3-4_plus. **E.** RMSD of Cα atoms during the simulation of EC3-4 monomer. The convergence of the trajectories was confirmed based on the stable RMSD values.

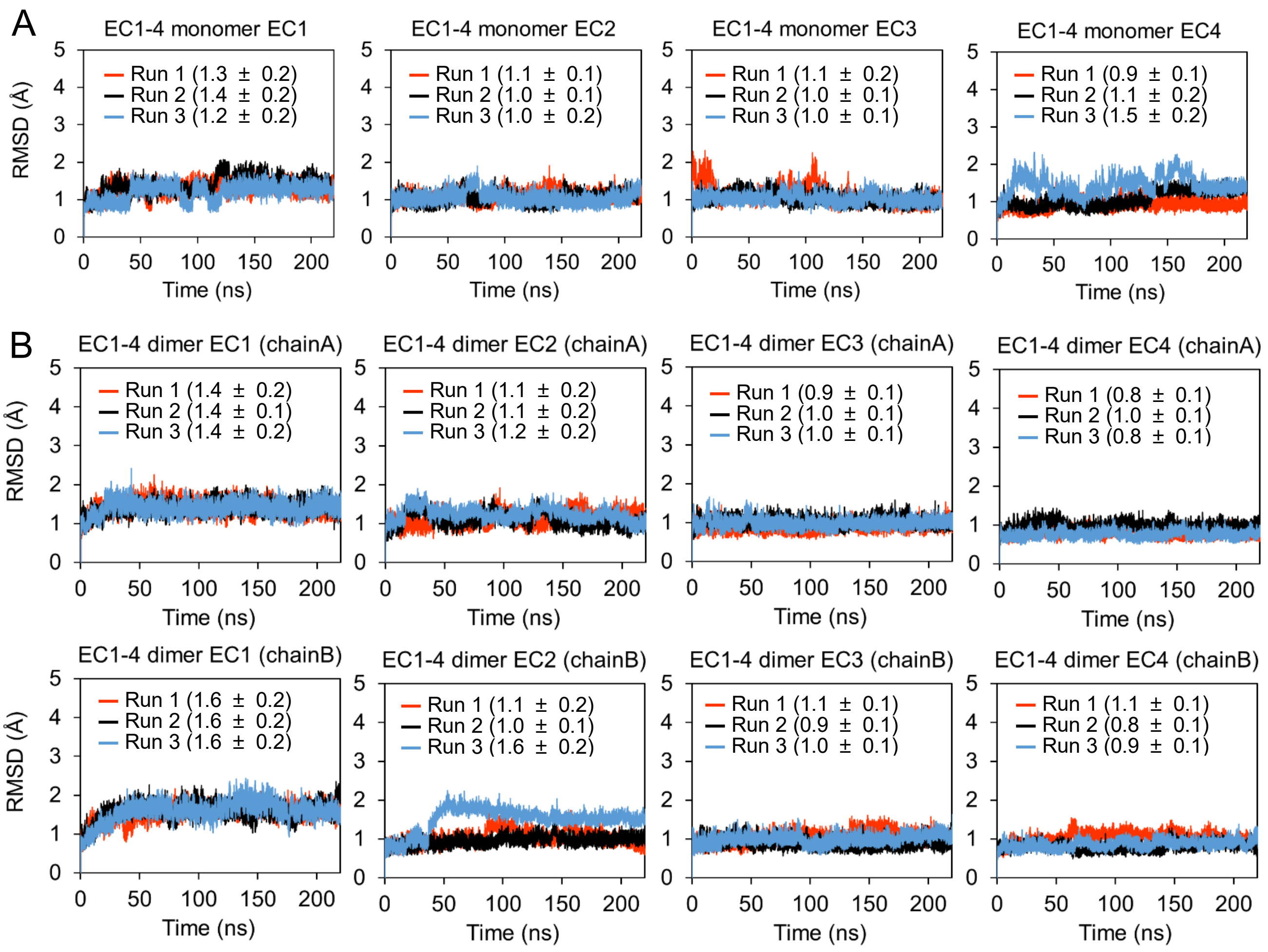


**Figure S2.** RMSD of Cα atoms during the simulations of EC1-4 WT monomer and homodimer. **A.** EC1-4 WT monomer, **B.** EC1-4 WT dimer. As the molecule showed high flexibility at $Ca^{2+}$-free linker, RMSD of each domain was calculated individually. Five Cα atoms at N-terminus were excluded from the calculation of RMSD of EC1 as they were disordered. Averages and standard deviations from 20 ns to 220 ns of each simulation are shown in the graph in angstrom units.

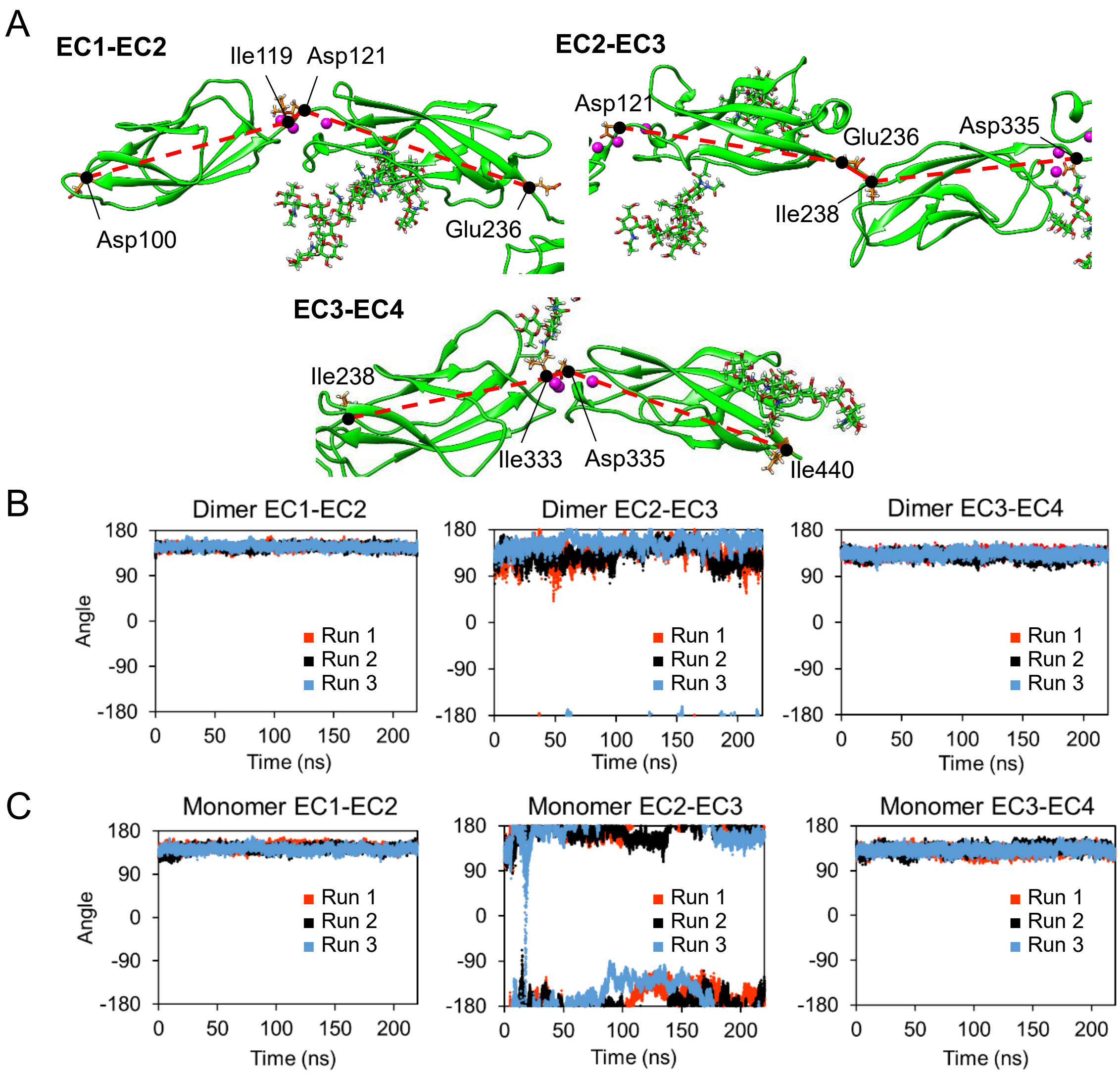


**Figure S3.** Dihedral angles of each domain. **A.** Atoms used for the calculation of dihedral angles. For the calculation of the angle between EC1 and EC2, the dihedral angle consisting of Asp100 Cα, Ile119 Cα, Asp121 Cα and Glu236 Cα was calculated. For the calculation of the angle between EC2 and EC3, the dihedral angle consisting of Asp121 Cα, Glu236 Cα, Ile238 Cα and Asp335 Cα was calculated. For the calculation of the angle between EC3 and EC4, the dihedral angle consisting of Ile238 Cα, Ile333 Cα, Asp335 Cα and Ile440 Cα was calculated. Cα atoms used for calculation are marked with black circle connected with red continuous and dotted lines. **B, C.** Dihedral angles between the adjacent domains of dimer (B) and monomer (C) calculated by GROMACS tools. The angles between EC2 and EC3 changes more drastically in the monomer.

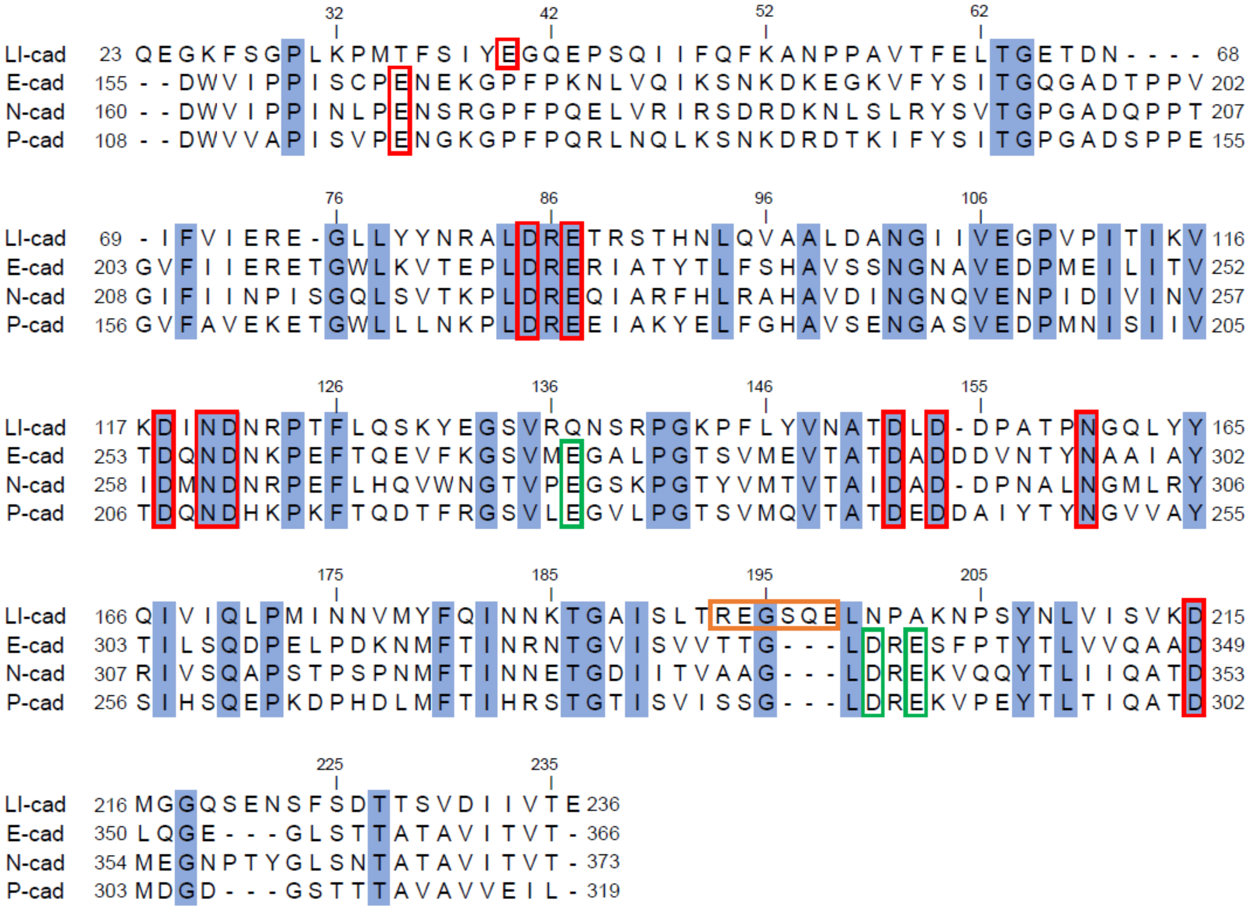

**Figure S4.** Sequence alignment of EC1-2 of human LI-, E-, N- and P-cadherin. Alignment was performed using CLUSTALW. Calcium binding motif for calcium ions between EC1 and EC2, and EC2 and EC3 are indicated by red and green boxes, respectively. LI-cadherin does not have binding motifs for calcium ions between EC2 and EC3. Residues forming α-helix in LI-cadherin EC2 are indicated by the orange box. Residues identical among the four cadherins are highlighted in blue.

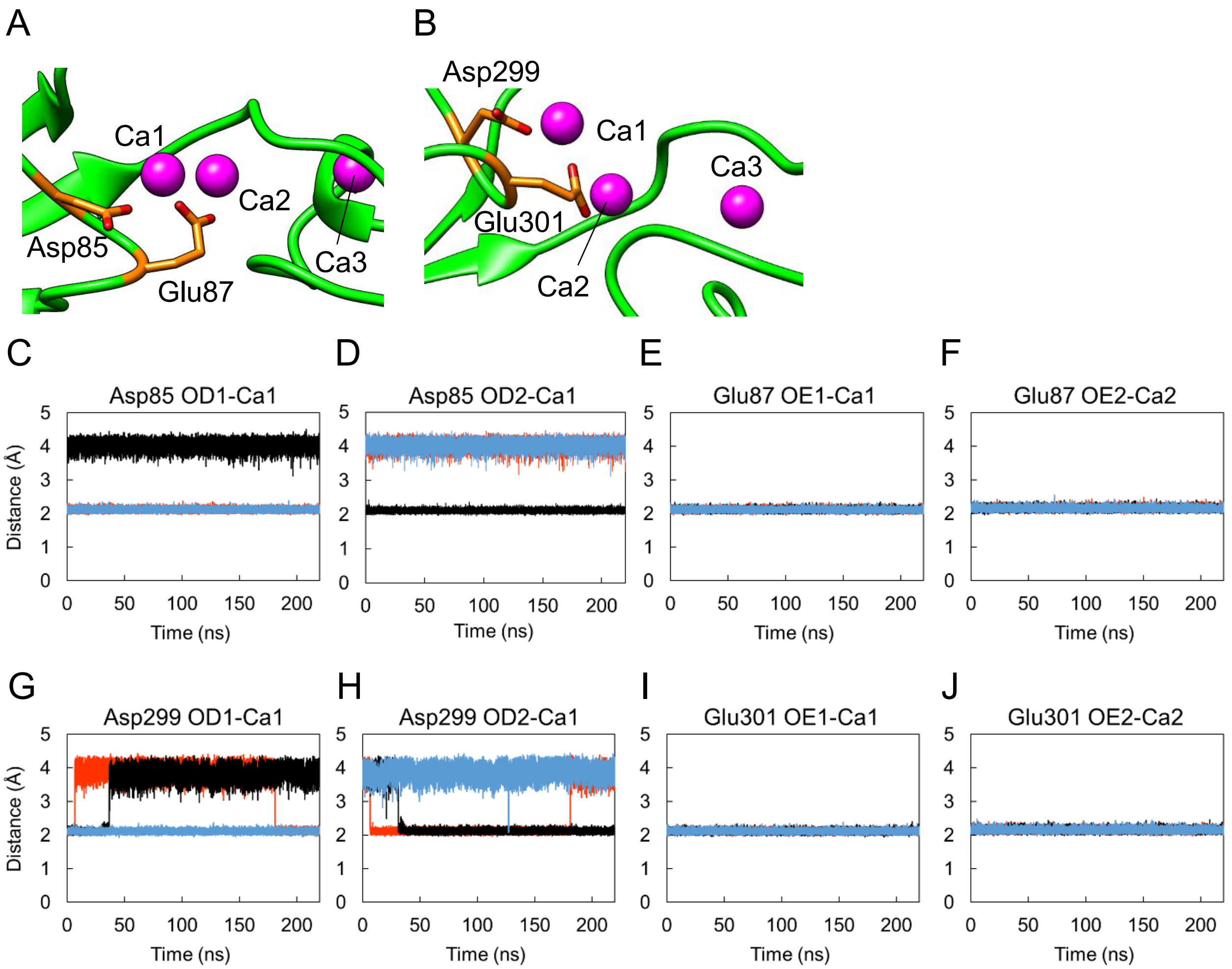


**Figure S5.** Coordination of Asp and Glu residues to Calcium ions. DXE motif in LI-cadherin EC1 (**A**) and EC3 (**B**). Residues belonging to DXE motif are shown in orange. **C~J.** Distances between Asp or Glu residue in DXE motif of EC1 or EC3 repeats and calcium ion 1 or 2. Distances between **C.** Asp85 OD1 and Ca2, **D.** Asp85 OD2 and Ca2, **E.** Glu87 OE1 and Ca2, **F.** Glu87 OE2 and Ca1, **G.** Asp299 OD1 and Ca2, **H.** Asp299 OD2 and Ca2, **I.** Glu301 OE1 and Ca2 and **J.** Glu301 OE2 and Ca1 are plotted against simulation time. Either OD1 or OD2 of Asp85 coordinated with Ca2 between EC1 and EC2. Glu87 OE1 and OE2 coordinated with Ca2 and Ca1 between EC1 and EC2, respectively. Either OD1 or OD2 of Asp299 coordinated with Ca2 between EC3 and EC4. Glu301 OE1 and OE2 coordinated with Ca2 and Ca1 between EC3 and EC4, respectively. Simulation was performed for three times. Each run is shown in red, black and blue.

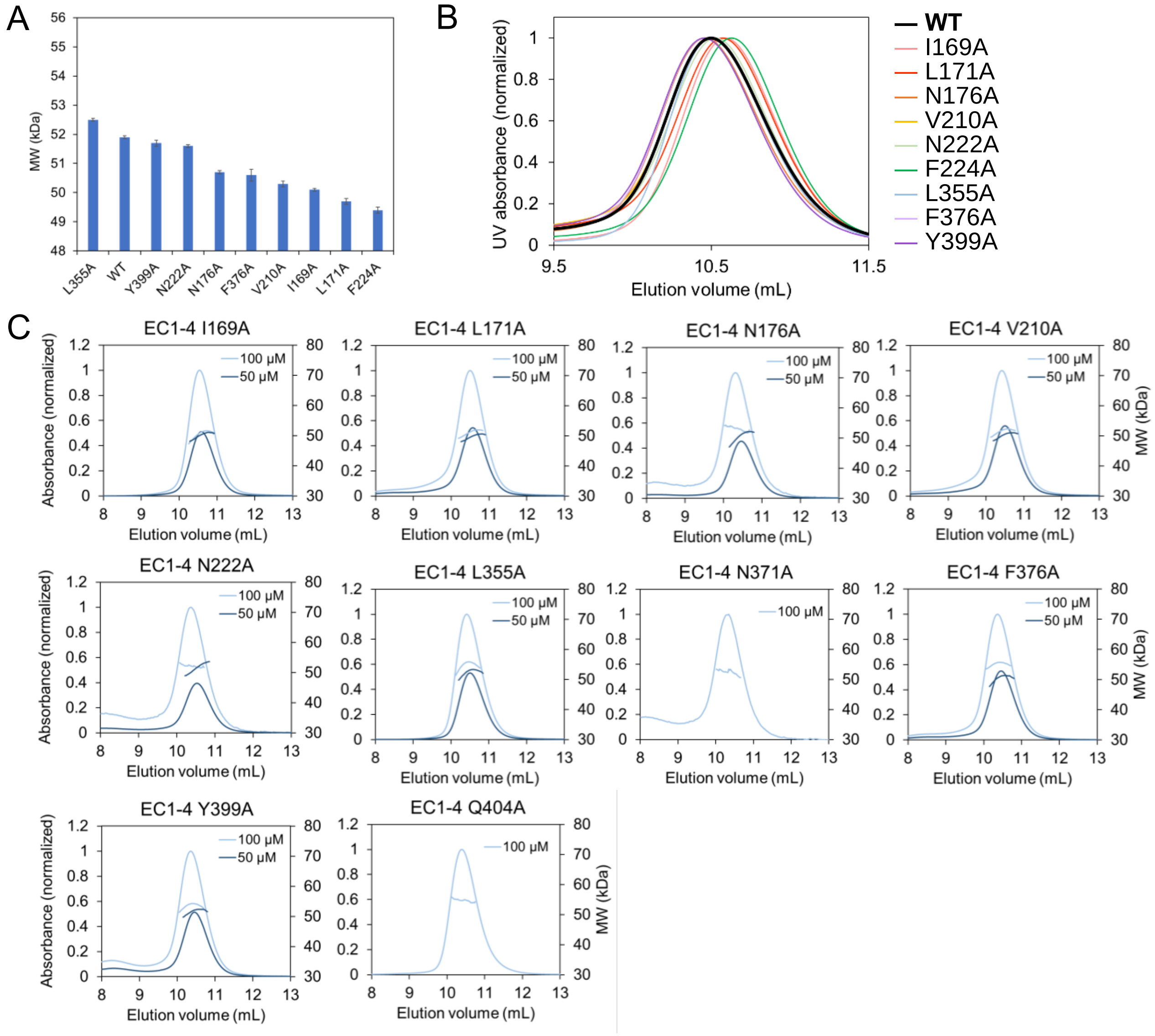


**Figure S6.** Results of SEC-MALS when the samples were injected at 50 µM. **A.** Molecular weight measured by MALS. Similar to the results when the samples were injected at 100 µM, F224A exhibited the smallest molecular weight among all constructs tested. **B.** SEC chromatograms obtained using SEC-MALS. Protein was injected at 50 µM. Chromatogram of wild type and F224A are indicated in black (bold line) and green, respectively. Elution volume of the peak top of F224A was the largest among all constructs. **C.** SEC-MALS results of EC1-4 mutants. Molecular weight measured by MALS is summarized in Table 2.

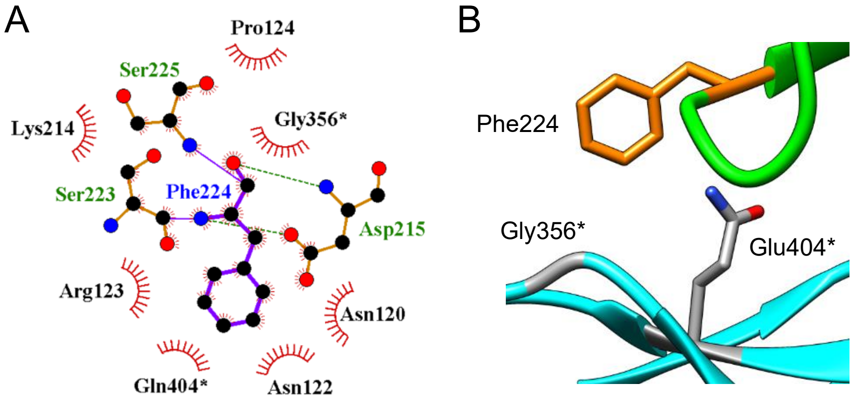


**Figure S7.** Residues interacting with Phe224 in chain A. **A.** Interaction depicted by LigPlot+ (64). Among the residues in chain B, only Gly356 and Gln404 were interacting with Phe224 in chain A. Residues belonging to chain B are indicated with an asterisk. **B.** Enlarged view of Phe224 in chain A (orange) and interacting residues in chain B (grey).

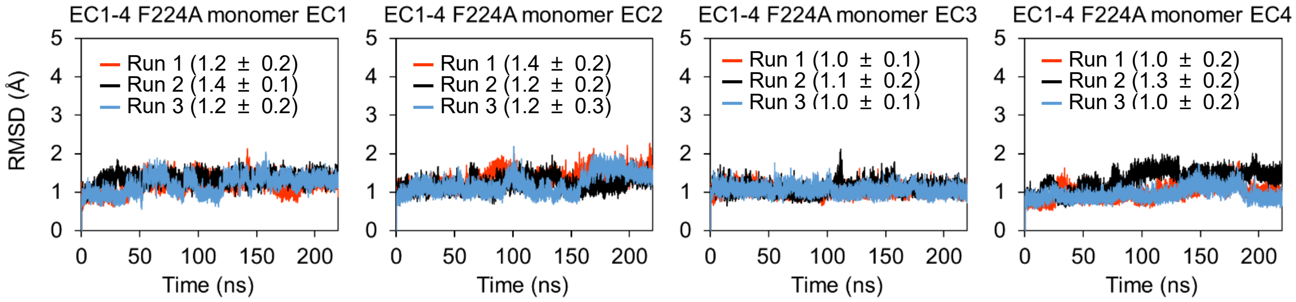


**Figure S8.** RMSD of Cα atoms during the simulations of EC1-4F224A monomer. As the molecule showed high flexibility at $Ca^{2+}$-free linker, RMSD of each domain was calculated individually. Five Cα atoms at N-terminus were excluded from the calculation of RMSD of EC1 as they were disordered. Averages and standard deviations from 20 ns to 220 ns of each simulation are shown in the graph in angstrom units.

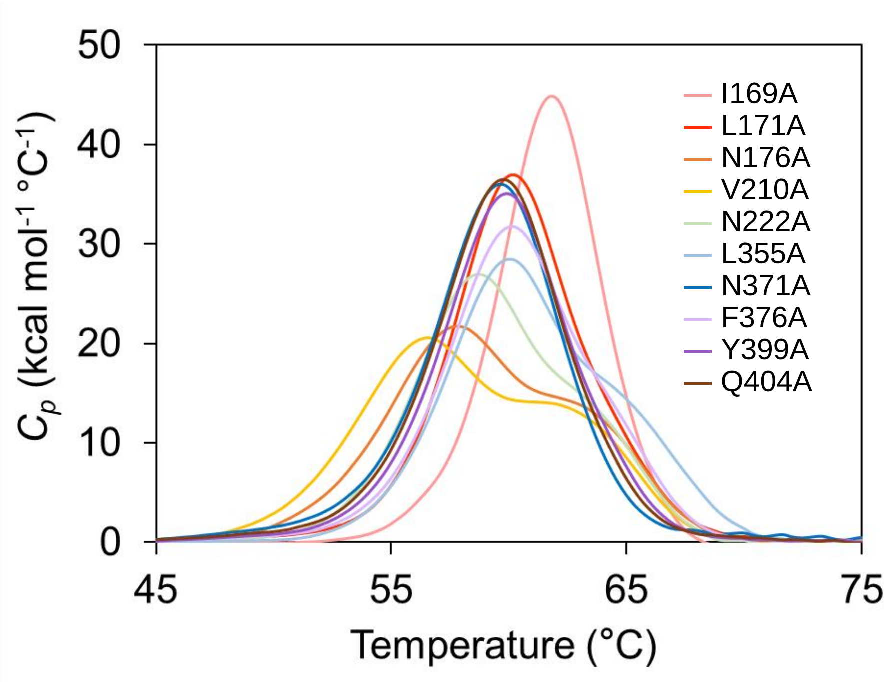


**Figure S9.** DSC data corresponding to EC1-4 mutants.

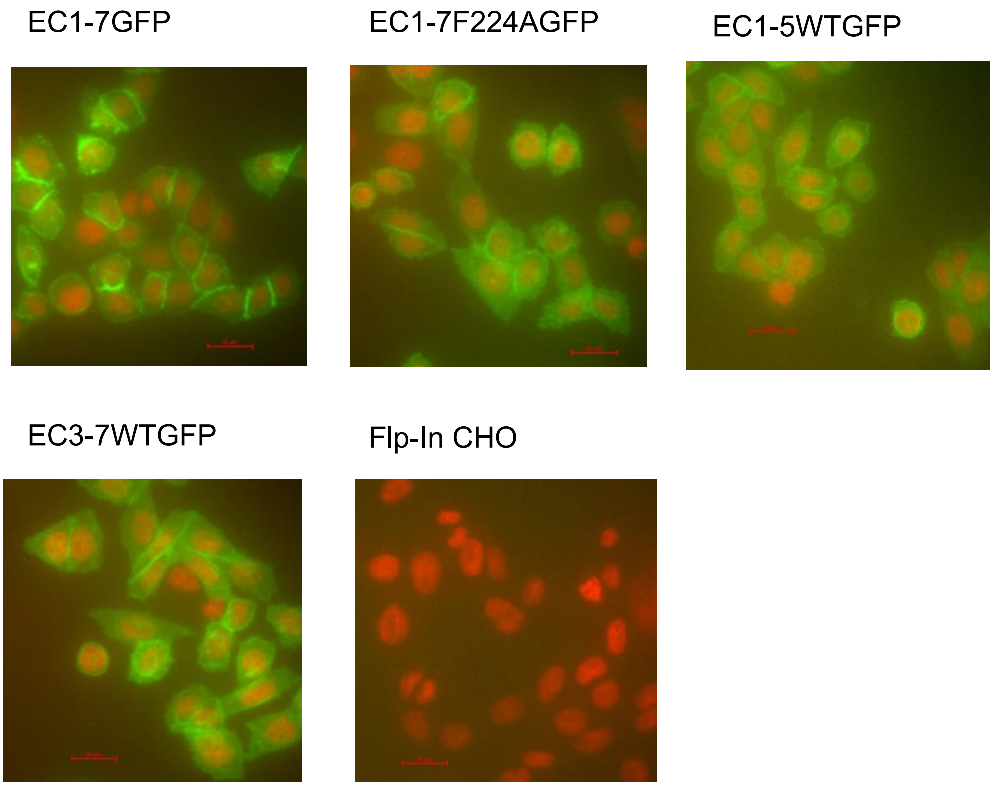


**Figure S10.** Verification of expression of LI-cadherin in mammalian cells. LI-cadherin expressing cells and mock cells were monitored with an In Cell Analyzer 2000 instrument. Fluorescence of GFP and Hoechst 33342 are shown in green and red, respectively. The constructs of LI-cadherin whose expression on the cell surface were verified are the following: EC1-7GFP, EC1-7F224AGFP, EC1-5GFP and EC3-7GFP. Non-transfected Flp-In CHO cells did not exhibit fluorescence derived from GFP. Scale bars (20 μm) are indicated with a red line.

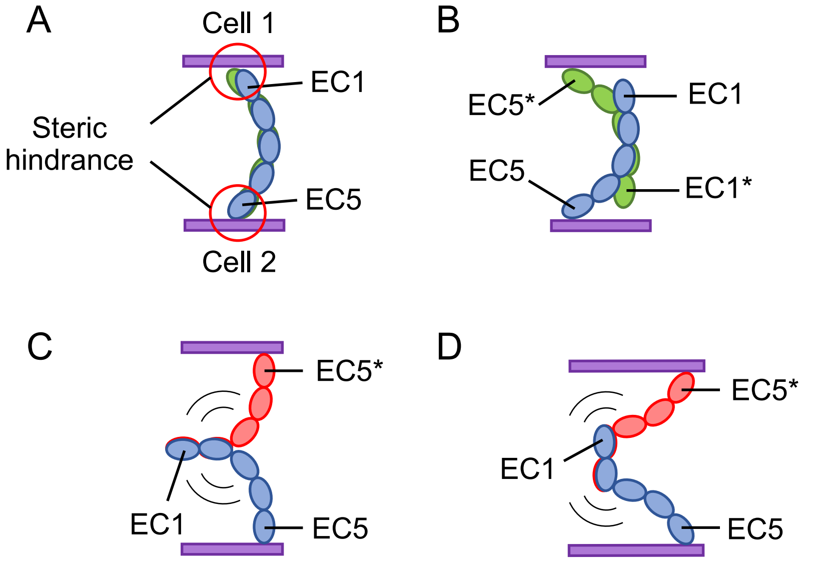


**Figure S11.** Schematic views of LI-cadherin EC1-5 homodimer. **A.** There seems to be steric hindrance between EC1 and cell membrane. **B.** EC2 and EC4 cannot interact with each other due to inappropriate orientation of LI-cadherin molecule. **C, D.** EC1-5 dimer formed by interaction of EC1-2 might not be formed due to instability of the molecules caused by high flexibility of $Ca^{2+}$-free linker between EC2 and EC3.

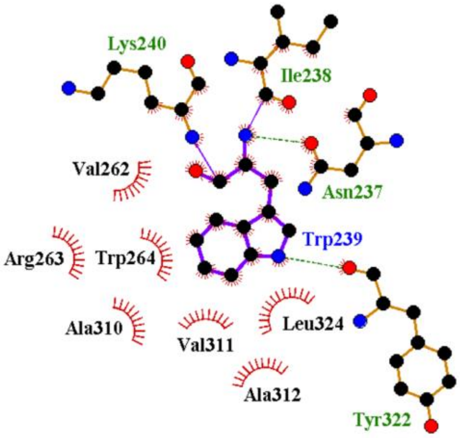


**Figure S12.** Residues interacting with Trp239 in chain A depicted by LigPlot+(64). Hydrophobic contacts are indicated in red lines. Hydrogen bonds are shown in green dotted lines.

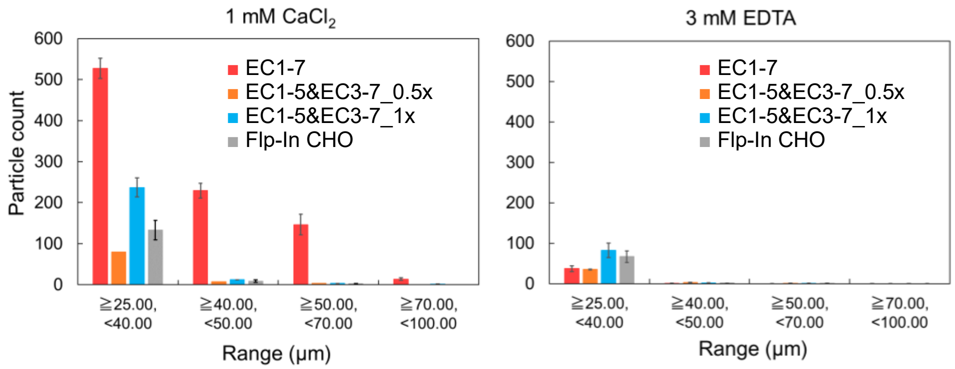


**Figure S13.** Size distribution of cell aggregates measured by MFI. Particles that were 25 μm or larger were regarded as cell aggregates. 1 x $10^5$ cells/mL solution of EC1-7 expressing cells and Flp-In CHO were used for the control measurements. For the measurement of the mixture of EC1-5 and EC3-7 expressing cells, 0.5 x $10^5$ cells/mL each (EC1-5&EC3-7_0.5x) or 1 x $10^5$ cells/mL each (EC1-5&EC3-7_1x) was used.

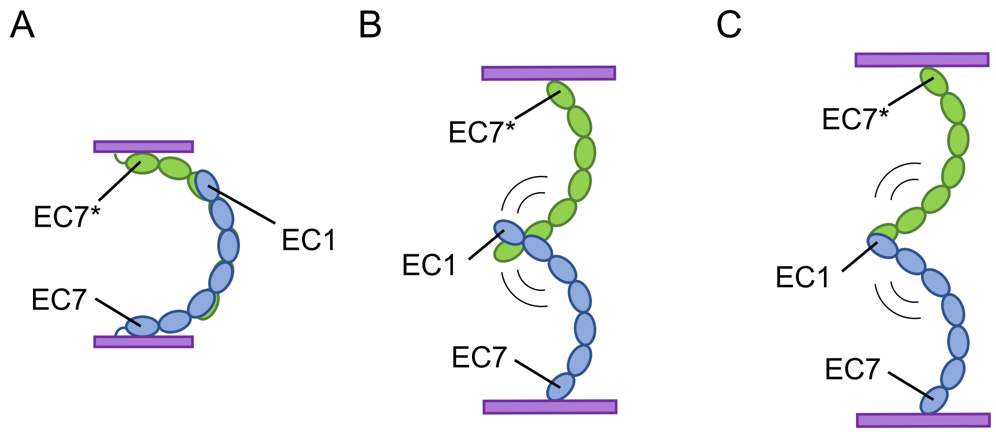


**Figure S14.** Schematic view of homodimer formed by LI-cadherin. **A.** EC1-4 homodimer observed by our crystal structure. **B.** X-dimer formed by EC1-2. **C.** Strand swap-dimer (ss-dimer) formed by EC1-2. As $Ca^{2+}$-free linker between EC2 and EC3 can move freely when X-dimer or ss-dimer was formed, these dimers seem not to be able to maintain LI-cadherin-dependent cell adhesion.
